# The Synthetic Epitope Atlas: High-Throughput Design and Validation of *De Novo* Antibody-Antigen Complexes

**DOI:** 10.64898/2026.04.17.719295

**Authors:** Nicholas Altieri, Joseph L. Harman, David Noble, Natasha Murakowska, Alexander Eng, Kerry L. McGowan, Davis Goodnight, Lucian DiPeso, Colleen Shikany, Emily Engelhart, Leah J. Homad, Miranda C. Lahman, Shyam Gandhi, Mackenzie Goodwin, Kendrick Herbst, Charles Lin, Margot McMurray, Juliana Barrett, Aditya A. Agarwal, James Harrang, Ryan O. Emerson, Randolph M. Lopez, David A. Younger, Adrian W. Lange

## Abstract

*De* novo antibody design models lack sufficient training data to reliably generalize. We demonstrate scalable generation of structural training data for machine learning-driven antibody design by linking *in silico* designs of antibody-antigen complexes to high-throughput experimental binding validation. Using AlphaSeq, a yeast-based platform for measuring protein binding affinities, we measure the affinity and specificity of thousands of *de novo* “synthetic epitope proteins” (SEPs) designed to bind to VHHs. The resulting Synthetic Epitope Atlas (SEPIA) pairs over 26 million on- and off-target affinity measurements with computationally designed VHH–SEP “pseudo-structures.” We validate strong, specific binding for 1,161 pseudo-structures and >75,000 VHH and SEP mutational variants. We show that these pseudo-structures complement existing structural databases and enable ML models to outperform confidence metrics commonly used to rank *de novo* antibody designs. Taken together, SEPIA establishes a scalable framework for improving *de novo* antibody design by augmenting sparse structural data with large-scale experimental binding data.

## 1 Introduction

Machine learning (ML) models are beginning to deliver on the promise of *de novo* design (DND) for antibodies, particularly for single-domain VHHs. Several groups have reported antibody DND campaigns with hit rates reaching half or more of attempted antigens, though results vary widely across studies and targets [1–10]. Critically, the ML models driving these advances [10–14] rely primarily on the Protein Data Bank (PDB) for training data, which contains approximately 250,000 experimentally determined protein structures [15, 16].

Despite much progress, antibody design pipelines generally yield low and inconsistent hit rates, and modern protein folding models struggle to accurately predict complexes for known antibody-antigen (Ab-Ag) pairs [14, 17–19]. This uneven performance suggests that current models learn some information about Ab-Ag complexes but lack sufficient scale and diversity of Ab-Ag training data. As of early 2026, only 10,763 PDB structures contain antibodies, including just 2,211 with VHHs but only 811 with unique VHHs in complex with their antigen [20, 21]. This scarcity of Ab-Ag structural data limits model improvement for antibody design, and the slow growth of the PDB indicates that traditional structural biology alone cannot resolve this bottleneck. Moreover, structural databases provide only positive binding examples, since non-binding pairs cannot form stable complexes for structure determination. As a result, datasets of solved structures lack informative “hard” negatives—plausible non-binding co-structures that are nonetheless essential for calibrating models and reducing false positives [22]. We believe that major advances in *de novo* antibody design hinge on radically expanding the scale and diversity of structural antibody training data.

To address these limitations, we generate large-scale training data by coupling DND of Ab-Ag complexes with high-throughput experimental binding validation using AlphaSeq [23], a yeast-based platform for measuring protein binding affinities that has been used across multiple studies to generate large-scale binding datasets and train ML models [24–29]. Prior work shows that experimentally validated *de novo* designed binders frequently and closely resemble their predicted binding geometries when verified by subsequent structural determination. [30–33] This enables predicted complexes to serve as structurally informative training examples when paired with quantitative binding measurements. We refer to these experimentally validated predicted complexes as “pseudo-structures,” which provide both positive examples (high-affinity binders that reinforce productive interaction geometries) and hard negatives (high-confidence designs that fail to bind) for model training.

To scale the generation of pseudo-structures for VHHs, we design synthetic epitope proteins (SEPs): *de novo* minibinders designed to bind VHH paratopes. By fixing the VHH “target” and designing diverse binders against it, we produce large numbers of predicted VHH–SEP complexes and experimentally measure binding using a high-throughput, standardized assay. Since minibinders historically validate at much higher rates than *de novo* VHHs, this strategy increases throughput while reducing experimental attrition [2, 4, 31, 34]. Moreover, since a single paratope can recognize multiple structurally distinct epitopes [35, 36], SEPs span diverse interaction geometries rather than collapsing onto a narrow motif class. This diversity provides rich training signal for ML, capturing what we refer to as an “epitope field”—i.e. the shared spatial distribution of biophysical properties across epitopes that bind a shared paratope.

In this study, we computationally design 45,430 SEPs against 190 unique VHHs across two design rounds and use AlphaSeq [23] to validate strong on-target binding for 1,161 VHH–SEP pseudo-structures. We additionally measure binding between thousands of VHH and SEP point mutants, generating more than 26 million on- and off-target binding affinity measurements and 76,470 VHH–SEP pairs that show strong on-target binding. We refer to this aggregate dataset as the Synthetic Epitope Atlas (SEPIA). We show that SEPIA pseudo-structures contain useful signal for ML training, particularly when combined with experimentally determined complexes from SAbDab-nano [20, 21]. Finally, we use SEPIA to train a binding classifier, ABACUS, that outperforms standard model confidence metrics on a holdout dataset of *de novo* designed VHH–Ag pairs, demonstrating that SEPIA data can improve the triage and prioritization of *de novo* antibody designs. Further details of our datasets, methods, experiments, and ML modeling are provided in the Supplementary Material.

## 2 Results

### 2.1 SEPIA Design Pipeline

To generate VHH–SEP pseudo-structures at scale, we developed a multi-stage structure-guided protein design pipeline (Figure 1A). In brief, starting from experimentally determined VHH structures (with cognate antigens removed), we generated *de novo* SEP backbones docked to VHH paratopes, designed sequences for each backbone, and applied structure-based filtering to retain candidates whose predicted VHH–SEP pseudo-structure closely matched the intended design geometry. High-confidence designs were then selected for experimental testing. Full pipeline details, including model configurations, filtering thresholds, and round-specific refinements, are provided in Supplementary Materials B and E. An analysis of the sequences and structures in SEPIA data is provided in Supplementary Materials C.

**Figure 1:**
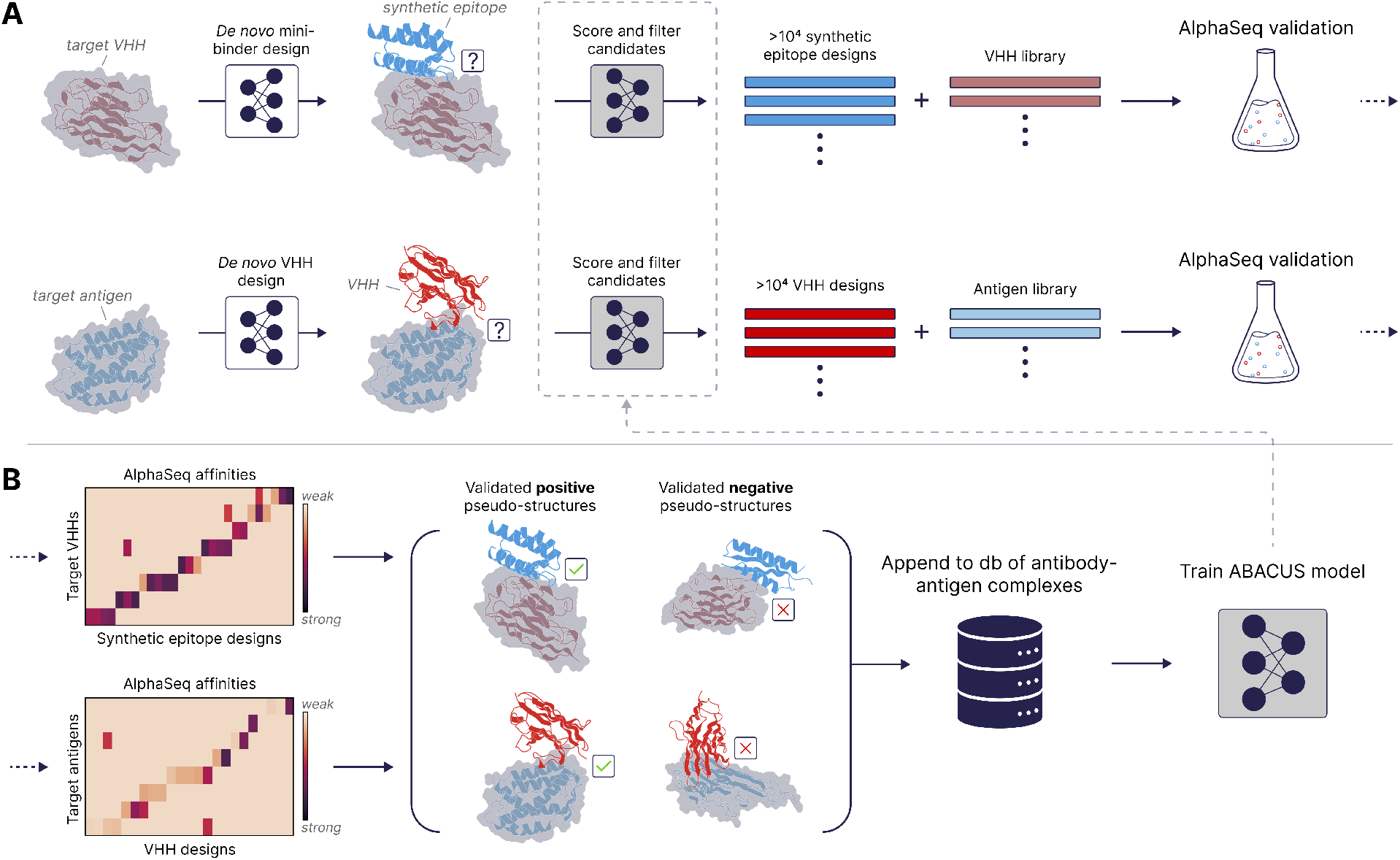
SEPIA workflow overview. Workflow for designing and validating synthetic epitope proteins (SEPs) and training ABACUS, our custom structure-based binding classifier. **(A)** *Design and measure:* Starting from a VHH structure, *de novo* SEPs are designed to bind to the VHH using a multi-stage structure-guided design pipeline, scored, and filtered to produce diverse candidate libraries (top). Analogously, *de novo* VHHs can be designed against antigen targets using a similar approach (bottom). Candidate binder and target libraries are assayed using AlphaSeq to obtain quantitative library-on-library affinity measurements. **(B)** *Validate and train:* AlphaSeq affinity measurements are used to identify validated “positive” pseudo-structures (strong, specific binding) and validated “negative” pseudo-structures (high-confidence designs that do not bind). These are appended to a database of antibody-antigen complexes and used to train our ABACUS model for filtering binders and non-binders.

SEPs were generated in two rounds. Round 1 targeted 48 VHHs and produced ∼600,000 candidate SEPs *in silico*, from which ∼25,000 were selected for AlphaSeq testing. Round 2 incorporated pipeline modifications and expanded the target set to 180 VHHs (38 repeated from Round 1 + 142 new), yielding ∼660,000 candidate SEPs and ∼20,000 experimentally tested SEPs. A third round of experiments focused on a select subset of Round 1 SEPs and VHHs to probe interface mutations. Together, these efforts established a scalable workflow for generating large libraries of predicted VHH–SEP complexes for high-throughput experimental validation and downstream ML modeling.

### 2.2 SEPIA *In Vitro* Validation Experiments

AlphaSeq binding experiments were performed for each SEPIA design round to experimentally validate predicted VHH–SEP pseudo-structures at scale. Binding interactions were categorized as a hit according to strong on-target affinity and specificity for the intended target (Supplementary Table S1).

#### 2.2.1 Round 1: Proof of Concept

In SEPIA Round 1, we screened 25,448 designed SEPs against 48 parent VHHs using AlphaSeq, generating more than 1.2 million pairwise protein–protein interaction (PPI) measurements (full heatmap shown in Figure 2A). Of these, 928 VHH–SEP interactions met hit criteria for strong, specific on-target binding (Figure 2B; Table 1; Supplementary Table S1), corresponding to 34 of the 48 VHHs (70.8%) and 3.6% of SEPs screened. Success rates varied widely across VHHs (Figure 2C), consistent with the variability observed in other DND studies [2, 6, 37]. Representative validated pseudo-structures for VHH 6U54 (Figure 2D) illustrate that diverse SEP designs engage distinct epitope geometries on the same VHH paratope, sampling the “epitope field” of shared biophysical properties recognized by a single paratope. Together, these results demonstrate that structure-guided DND of SEPs against VHHs can reliably produce high-affinity, highly specific, and diverse VHH–SEP pseudo-structures. Additional analyses of SEP physicochemical properties, structural diversity, and PDB novelty are provided in Supplementary Material C.

**Table 1.** SEPIA designs tested against wild-type VHHs, used for ML model training.

| Dataset | VHHs | SEPs | VHH Hits (%) | Validated VHH-SEP Positive Pseudo-structures (%) |
| --- | --- | --- | --- | --- |
| Round 1 | 48 | 25,448 | 34 (70.8%) | 928 (3.6%) |
| Round 2 | 180<sup>a</sup> | 19,982 | 75 (41.7%)<sup>b</sup> | 233 (1.2%) |

<sup>a</sup>142 new + 38 from Round 1. <sup>b</sup>25 from Round 1 + 50 new.

**Figure 2:**
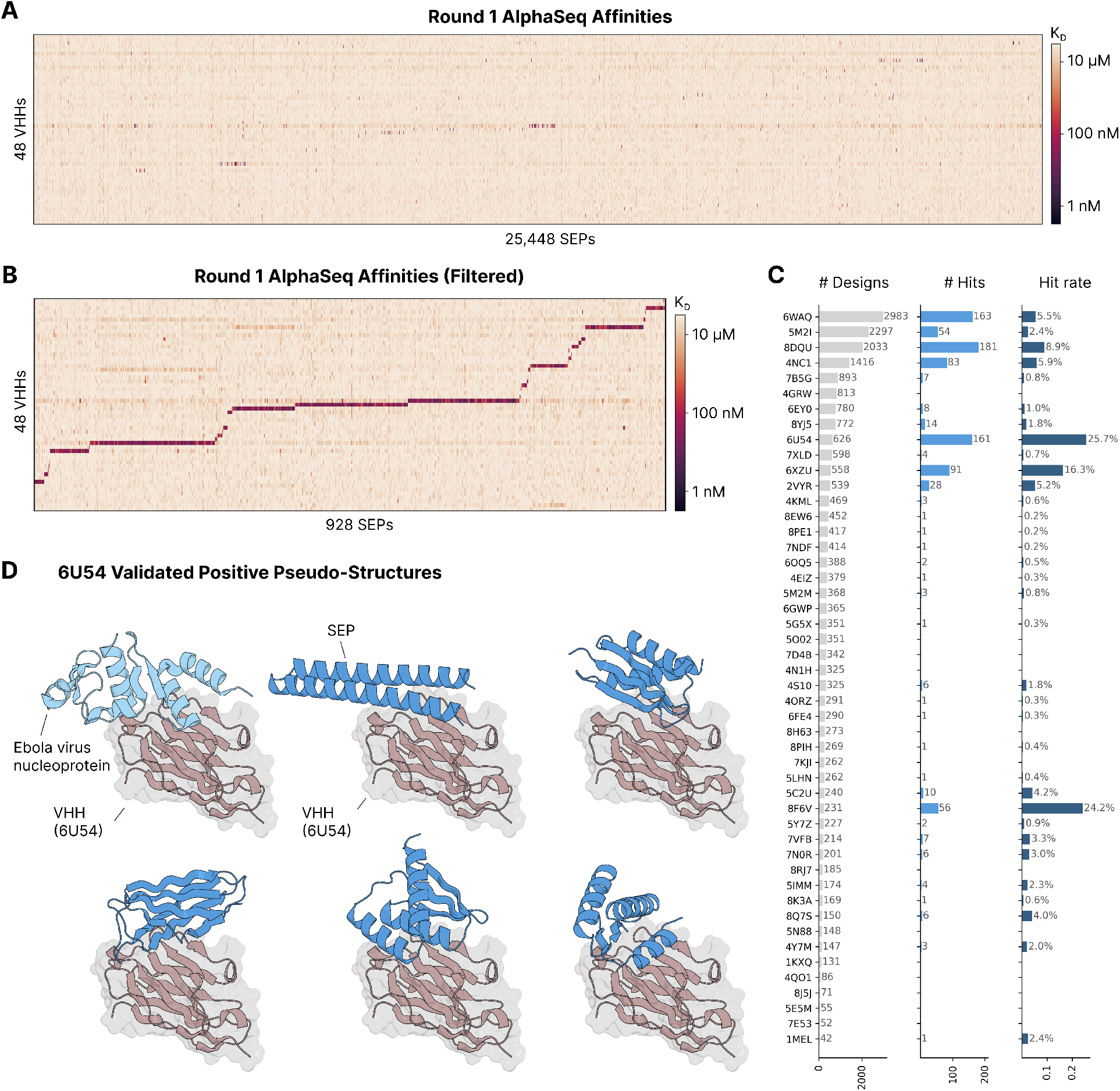
Round 1 AlphaSeq validation results. **(A)** AlphaSeq affinity heatmap for 25,448 SEP designs screened against 48 VHHs. Color scale indicates AlphaSeq binding affinity strength (darker = stronger binding). **(B)** Filtered heatmap showing the 928 SEPs meeting hit criteria, sorted by the VHH they bind. **(C)** Number of SEP designs tested per VHH (left), number of SEP hits per VHH (center), and hit rate per VHH (right), illustrating variability of design success across VHH targets. **(D)** Representative validated pseudo-structures for VHH 6U54. The experimentally determined complex of a VHH bound to the Ebola virus nucleoprotein (PDB: 6U54) is shown first (VHH in red, antigen in light blue), followed by designed VHH 6U54–SEP pseudo-structures (VHH in red, SEPs in dark blue). SEPs engage distinct epitope geometries on the same VHH paratope.

#### 2.2.2 Round 2: Paratope Local Diversity

Building on the success in Round 1, we expanded the diversity of VHH paratopes represented in SEPIA to increase structural coverage, test the generalizability of the design pipeline, and broaden the training data available for down-stream ML modeling. Round 2 targeted 180 parent VHHs, including 38 VHHs previously tested in Round 1 and 142 newly introduced VHHs to expand paratope diversity (Table 1).

In total, we screened 19,982 SEPs against 11,425 total VHHs (180 parent VHHs + ∼ 50–100 point mutants of each), generating over 26 million PPI measurements (Supplementary Figure S15A). Round 2 produced 233 parent VHH–SEP pseudo-structures meeting hit criteria, corresponding to 75 of 180 VHHs (41.7% hit rate, including 25 successfully targeted VHHs from Round 1 and 50 new VHHs) and representing 1.2% of SEPs tested (Table 1). The reduced hit rates in Round 2 likely reflect the broader VHH space explored, equal allocation of top-ranked SEPs per VHH, and pipeline modifications (notably Boltz-2 “hard templating” [14, 38] and a custom RFDiffusion potential based on 3Di structure encoding [39]) that were later found to impact hit rate and binder discrimination. Despite reduced hit rates, the absolute number of VHHs with at least one validated SEP nearly doubled, expanding VHH diversity in the aggregate SEPIA dataset.

As in Round 1, binding was dominated by strong on-target interactions with limited off-target activity, indicating that specificity was largely preserved despite increased SEP and VHH diversity. To probe paratope local sequence sensitivity, Round 2 additionally included 11,425 single-point mutant VHHs. Across these mutant libraries, we observed 36,803 hit interactions involving 3,855 mutant VHHs (33.7%) and 1,435 SEPs (6.9%) (Table 2).

**Table 2.** SEPIA mutational datasets for probing the local mutational landscapes of SEP-VHH pseudo-structures.

| Dataset | Mutation Type | Parental |  | Variant |  | Hits (%) |  | Validated VHH-SEP Mutant Hits (%) |
| --- | --- | --- | --- | --- | --- | --- | --- | --- |
|  |  | VHH | SEP | VHH | SEP | VHH | SEP |  |
| Round 2 | VHH point | 180 | 20,816<sup>a</sup> | 11,425 | – | 3,855 (33.7%) | 1,435 (6.9%) | 36,803 (2.7%) |
| Round 3 | SEP SSM | 1 | 1 | – | 4,779 | 1 (100%) | 3,496 (73.2%) | 3,496 (73.2%) |
| Round 3 | VHH+SEP ala | 4 | 96 | 57 | 7,040 | 56 (98.2%) | 5,488 (78.0%) | 35,010 (30.7%) |

<sup>a</sup>Includes Round 1 SEP hits for testing against VHH variants.

Analysis of VHH point mutations reveals that affinity-sensitive residues differ between SEPs targeting the same VHH, consistent with each SEP engaging a distinct subset of paratope residues. These residues localize to the predicted binding interface (Supplementary Figure S15; Supplementary Material C.4.1), revealing a rich landscape of tolerated and deleterious mutations at the VHH–SEP interfaces. Together, these data substantially increase the scale and granularity of joint sequence–structure–affinity data within SEPIA.

#### 2.2.3 Round 3: Interface Validation via Mutational Scans

SEPIA Round 3 complements Rounds 1 and 2 by systematically probing the sequence determinants of VHH–SEP binding through targeted mutational scans. This round comprises two complementary experiment types (Table 2).

##### SEP single-site mutagenesis

To identify SEP residues critical for binding, we performed site saturation mutagenesis (SSM) on validated SEP designs targeting a single VHH (7XLD), yielding 4,779 SEP variants total (single-site substitutions at every position plus randomly selected double mutants). Of these, 3,496 variants (73.2%) retained strong binding, enabling position-resolved mapping of mutational tolerance across the SEP sequence. Mutations at SEP residues in contact with the VHH in the designed pseudo-structure tend to be most disruptive to binding affinity (Figure 3 and Supplementary Materials C.4.3).

**Figure 3:**
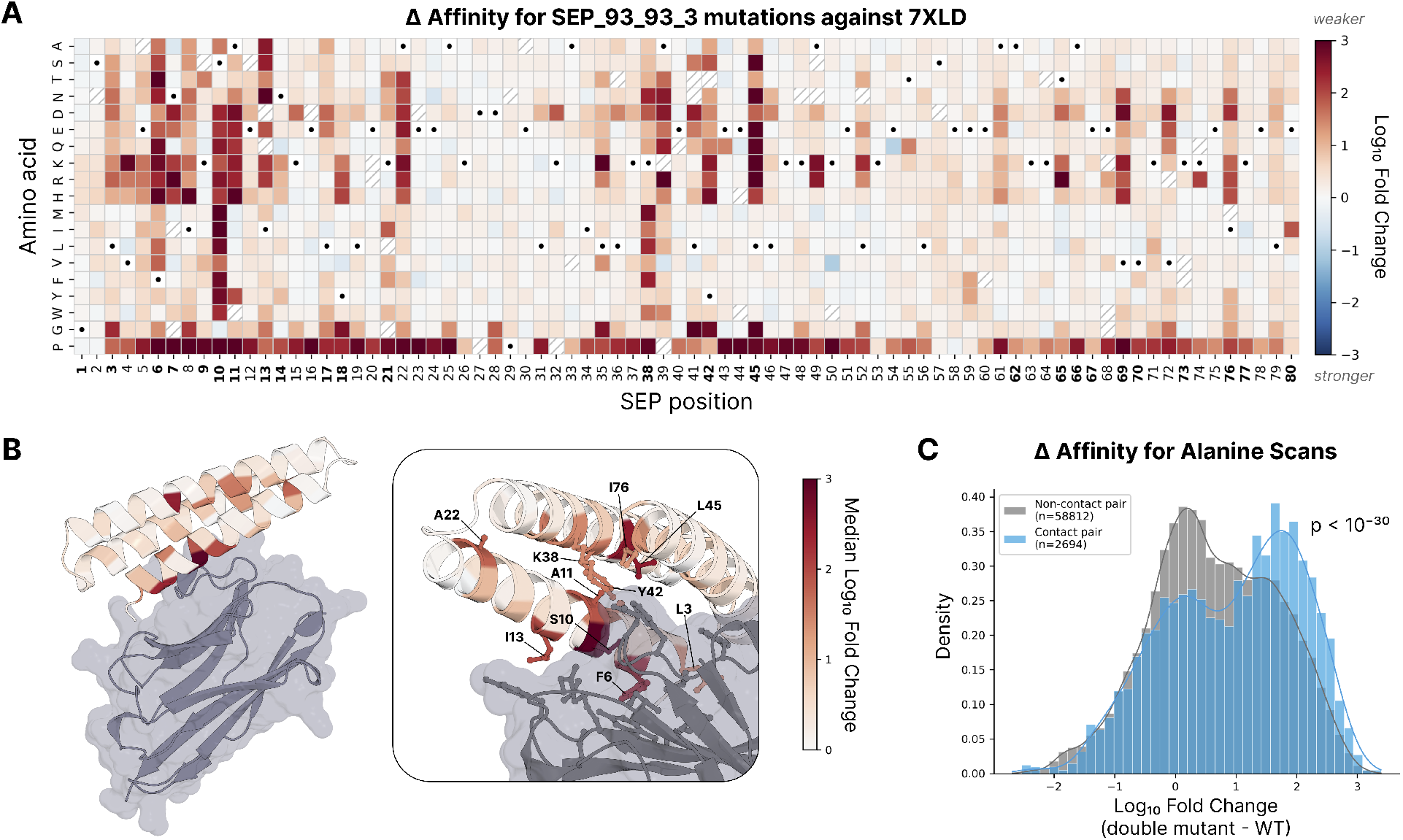
Round 3 interface validation results. **(A)** SSM results showing the change in binding affinity for each mutant of a SEP binding to VHH 7XLD. Black circles indicate the parental SEP amino acid identity at each position. Diagonal hatching marks mutations that failed to express in AlphaSeq. Residue positions likely involved directly in binding are shown in bold on the x-axis, defined as SEP residues with any heavy atom within 4.5 Å of any VHH heavy atom in the pseudo-structure. **(B)** Pseudo-structure of the VHH 7XLD–SEP complex, displayed at full scale (left) and with an interface close-up (right). The SEP is colored according to the median affinity change per residue (excluding mutations to proline), while VHH 7XLD is shown in gray. In the interface view (right), the 10 SEP residues exhibiting the largest median affinity changes are labeled with side chain heavy atoms displayed. Contacting VHH side chains are also shown, illustrating how the CDR3 loop of VHH 7XLD penetrates the SEP core region where mutagenesis causes the greatest binding disruption. **(C)** Distributions for change in affinity of alanine scan mutations at non-contact residues (gray) versus contact residues (blue) for 96 Round 3 VHH–SEP pseudo-structures. The contact residue distribution is significantly shifted toward higher fold change, indicating that mutants at the VHH–SEP interface are more likely to disrupt binding.

##### Double-sided alanine scanning

To validate predicted VHH–SEP contact interfaces, we systematically mutated each amino acid position to alanine on both the VHH and SEP sides. For each VHH, we generated CDR3 alanine-scan variants of the VHH (57 total unique VHH variants) and alanine-scan variants of 24 cognate SEPs each (7,040 total unique SEP variants). This double-sided design enables direct comparison of mutational effects on both binding partners. Residues predicted *in silico* to lie at the VHH–SEP interface exhibited significantly greater affinity ablation upon alanine substitution compared to non-interface residues (Mann-Whitney *U* test, *p <* 10^−30^; Figure 3C), with residual effects at non-interface positions attributable to impacts on protein folding stability [24]. This function-to-structure concordance, in addition the SSM results, provides further validation that designed SEPs engage their target VHHs through the intended structural interfaces, lending experimental support to the computational models underlying SEPIA design.

### 2.3 SEPIA Model Evaluation Tasks

We constructed two evaluation tasks to assess models trained on SEPIA data.

#### 2.3.1 SAbDab-nano Decoy Dataset

To benchmark generalization on native VHH–antigen complexes, we constructed a large dataset of known binders and AlphaSeq-validated non-binders (decoys) from SAbDab-nano. For on-target interactions (known binding pairs), we used the known PDB co-structures as templates. For off-target interactions (AlphaSeq-validated non-binders), we employed a two-stage prediction process. In Stage 1, we generated Boltz-2 predictions templated with their respective AlphaFold-predicted antigen monomer structures as initial complex predictions. In Stage 2, we re-predicted each VHH-antigen pair using the best Stage 1 complex prediction as a template, ensuring comparable templating between on-target and off-target sets. Three predictions were generated per interaction without multiple sequence alignments (MSAs); the prediction with the highest ipSAE (interaction prediction Score from Aligned Errors) score [40, 41] was selected for each PPI. The final benchmark dataset included 15,167 unique PPIs (646 on-target, 14,521 off-target) spanning 960 unique VHH and 438 unique antigens, providing a 22.5:1 decoy ratio. On-target predictions and decoys were labeled “bind” and “non-bind,” respectively. The resulting dataset is akin to that of Smorodina et al. [42], which defines a classification task: distinguishing true binders from structurally plausible non-binders (decoys). This task is analogous to filtering high-confidence from low-confidence designs in DND pipelines.

#### 2.3.2 *De Novo* VHH Dataset

A key question is whether a classifier trained on SEPIA’s synthetic VHH–SEP pseudo-structures can generalize to real *de novo* antibody design campaigns, where VHHs bind native antigens rather than engineered epitopes. To test this, we used several open-source *de novo* VHH design models to generate and experimentally test 4,930 candidate VHHs against 22 antigens, none of which have an antibody-bound structure in the PDB. AlphaSeq validation identified 37 high-quality hits across 8 antigens (36.4% antigen hit rate, 0.8% design hit rate). These experimentally validated designs, together with their predicted *in silico* structures, serve as hold-out data for a stringent test of model generalization to DND campaigns.

Structural analysis via Foldseek [39] revealed that validated *de novo* VHH–antigen complexes show minimal structural similarity to the PDB (median complex TM-score 0.455), confirming that these designs explore novel structural space. Furthermore, SEPIA VHH–SEP complexes and *de novo* VHH-antigen complexes form entirely distinct structural clusters at both the complex and monomer levels, with zero mixed clusters observed. This indicates that designed synthetic epitopes adopt structural modes distinct from native antigen binding interfaces. Additional details are provided in Supplementary Material D.2.

### 2.4 SEPIA Model Evaluation Results

We conducted a set of *in silico* experiments to evaluate the information content of SEPIA data. Experiments described in this section make use of our Neighborhood Transformer (NTX) model (Supplementary Material E.2). To control for information leakage and fairly evaluate performance on the decoy dataset, NTX is pre-trained on only monomer protein structures from AFDB [43], excluding any antibodies in the PDB, such that its embeddings are blind to Ab-Ag binding.

#### SEPIA pseudo-structures encode learnable binding signal

To examine whether *in vitro* validation provides meaningful signal for ML beyond simply training on *in silico* designs, we performed a label ablation study in which *in vitro* bind/no-bind labels were randomly permuted in a 5-fold cross-validation and a model was tasked with predicting held-out labels. We concluded at a *p*-value < 0.001 that distinguishing binders from non-binders in a cross-validation experiment is possible, but only when training on the real *in vitro* validation labels and only with a representation aware of the Ab-Ag complex structure. Results are shown in Supplementary Figure S20. A complementary analysis (Supplementary Figure S21) showed that AlphaSeq-derived labels are more informative than pseudo-labels based on the confidence metrics ipSAE.

#### Model performance tends to improve with more SEPIA data

To assess the relative value and interoperability of SEPIA pseudo-structures with experimentally-derived co-structures, we built simple models to perform the decoy detection task using varying numbers of real SAbDab-nano structures, with or without supplementation with SEPIA pseudo-structures. Briefly, we trained a simple multi-layer perceptron (MLP) neural network classifier on 100 stratified resamplings of our 5-fold cross-validation training sets (Supplementary Material F.1) across a range of data volume. The SAbDab-nano decoy data, containing a total of 646 real VHH-Ag complexes, is effectively capped at sampling 400 positive structures due to the cross-validation splits. SEPIA is similarly capped at 750 positive pseudo-structures. The performance of MLP classifiers on only SAbDab-nano data was compared to training on the same SAbDab-nano data plus SEPIA data to produce an estimate of relative informational value.

SEPIA data is ostensibly out-of-distribution for this task (having SEPs instead of natural antigens), yet, as Figure 4 demonstrates, SEPIA data is complementary to the PDB-derived decoy data and lifts model performance at detecting real VHH–Ag complexes versus decoys. While the learning curve for training on only the decoy dataset is capped by available PDB data, the addition of SEPIA data allows the learning curve to continue its upward trend, demonstrating that pseudo-structures are interoperable with real experimentally determined co-structures and have comparable signal for the purposes of model training. Further details of the learning curve methodology are provided in Supplementary Material F.4.

**Figure 4:**
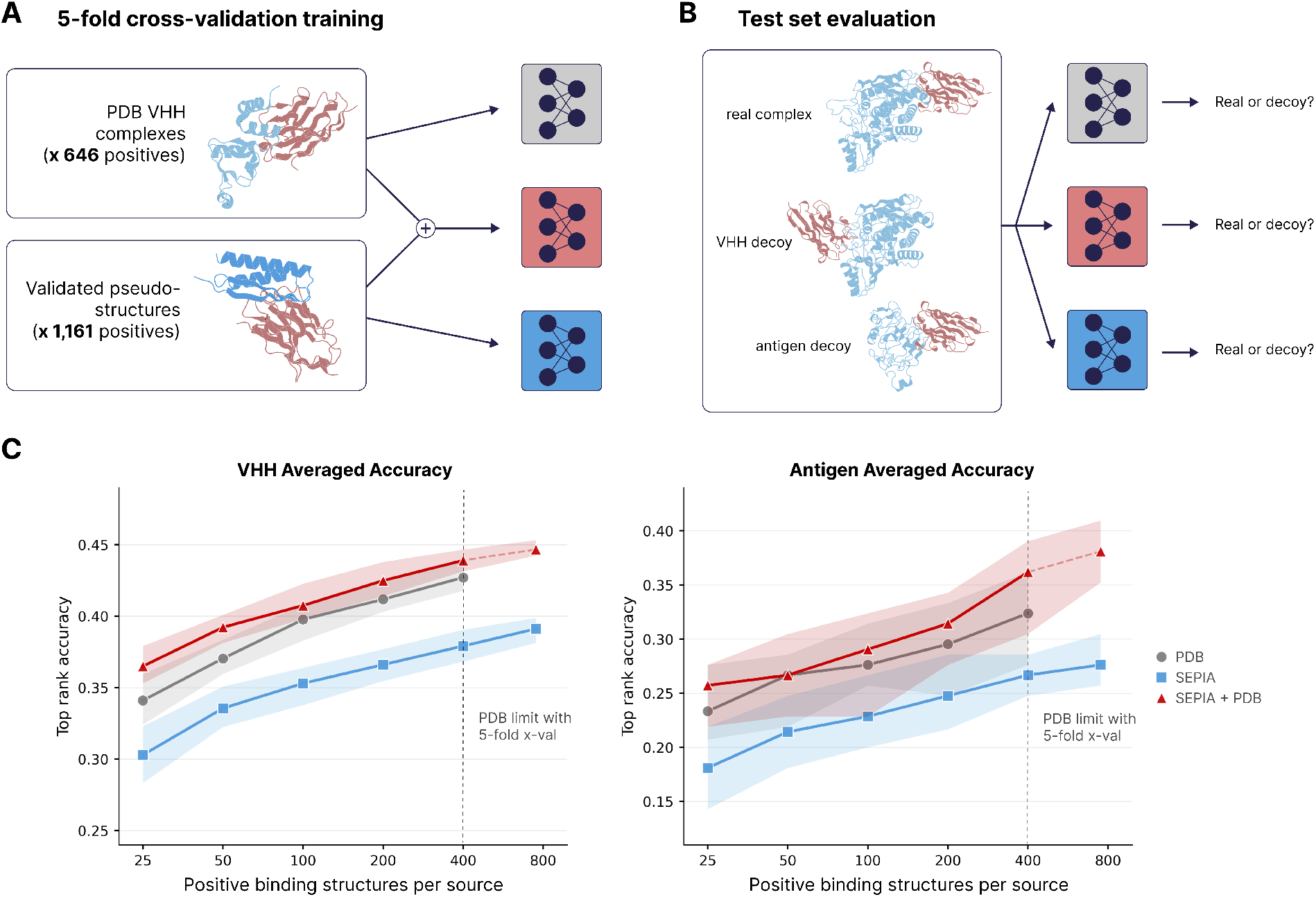
Cross-validation learning curves. **(A)** The decoy dataset (comprising real VHH-antigen complexes from the PDB) and the SEPIA dataset are used as training data within a 5-fold cross-validation experiment. Simple MLP classifiers are trained on resamplings of the PDB-derived decoy dataset (gray), the SEPIA dataset (blue), and the union of both (red). **(B)** Trained MLPs are evaluated on their corresponding decoy data test set split at the task of predicting if a VHH-antigen complex is a real or decoy structure. Decoys consist of either an antigen with a mismatched VHH or a VHH with a mismatched antigen. **(C)** Model performance at varying training data volumes is evaluated as top rank accuracy: the fraction of cases in which the true VHH–antigen complex ranks higher than all decoys for subsets containing the same VHH (left) or containing the same antigen (right). Accuracy is computed only for subsets with ≥2 decoys, and averages are taken over the entire decoy set by concatenating the predictions on all 5 test sets within each cross-validation pass. Curves represent the median (error bars demarcate the 25th and 75th percentiles) from the 100× resamplings of the training sets at a 1:10 ratio of binder:non-binder structures. The baseline accuracy averaged by VHH and by antigen is 0.065 and 0.024, respectively.

### 2.5 Increasing DND Efficiency with SEPIA

To evaluate the utility of pseudo-structures on real design tasks, we assessed the ability of a model trained on SEPIA data to enrich for binders in a recent DND campaign (Section 2.3.2; Supplementary Material D.2). Candidate filtering is a key step in most modern DND pipelines, and ranking algorithms are fundamentally modular and relatively model-agnostic. We therefore trained a deep learning-based classifier (built on top of Boltz-2 [14]) on SEPIA data to predict *in vitro* binding from proposed co-structure.

#### 2.5.1 ABACUS

We introduce ABACUS (**A**nti**B**ody **A**ffinity **C**lassifier **U**sing p**S**eudo-structures), our classifier model for filtering *de novo* VHH–Ag designs (Figure 5).

**Figure 5:**
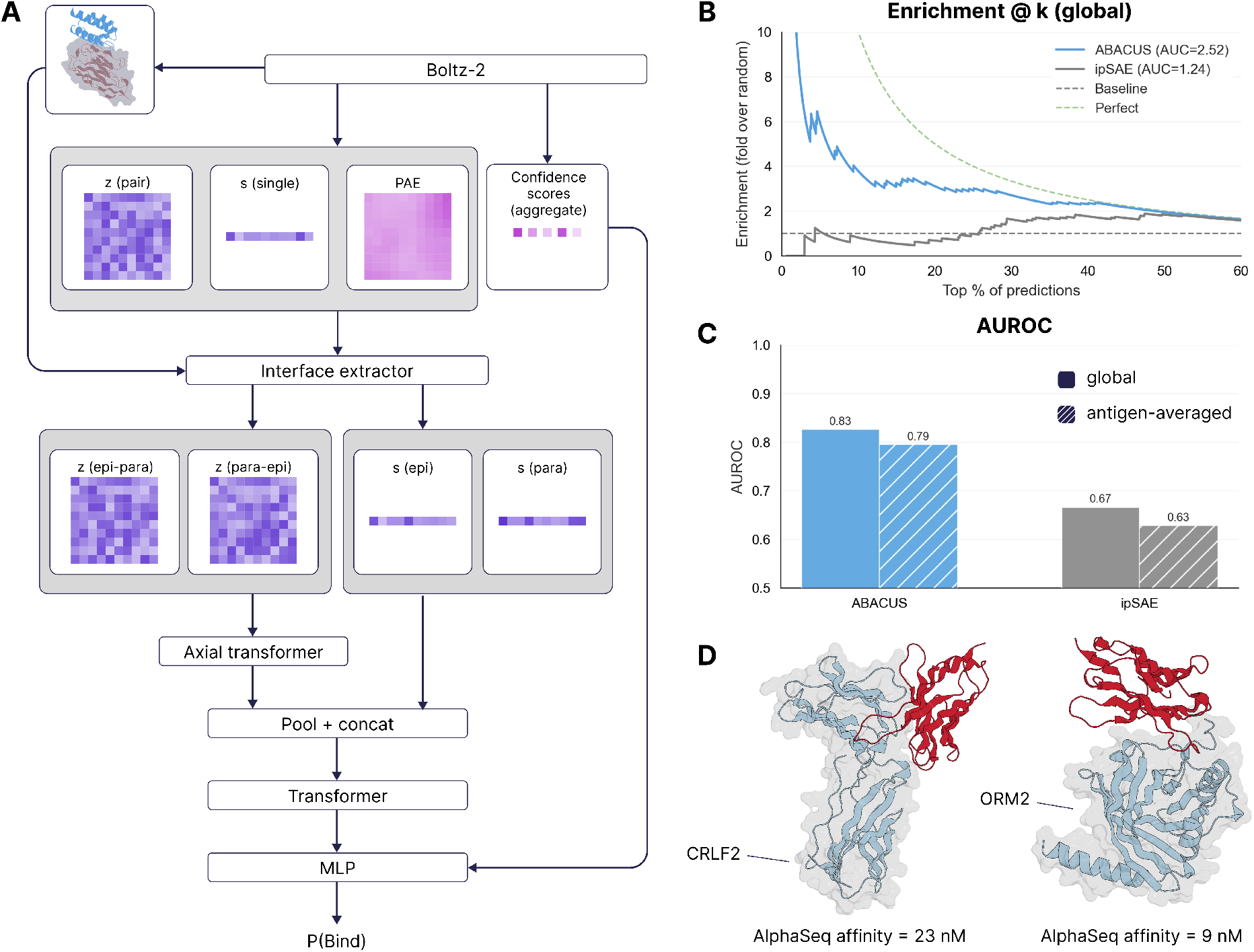
The ABACUS model and its performance on the *de novo* VHH dataset. **(A)** ABACUS model architecture overview: Boltz-2 is used to generate structure predictions, embeddings, and confidence scores. The predicted structure defines the Ab–Ag interface used to aggregate embeddings over residues on the epitope and paratope, which then feed into a series of transformer and pooling layers before finally outputting a classification prediction for binding. **(B)** Enrichment@k on the held out *de novo* VHH dataset, evaluated globally over all antigens. The confidence metric ipSAE (gray) displays worse-than-random enrichment at the highest percentiles, while ABACUS (blue)—trained on the union of SEPIA and decoy datasets—displays enrichment ≥4 for the top 10% of designs. **(C)** AUROC on the held out *de novo* VHH dataset, evaluated globally and averaged by each antigen with at least one hit. ABACUS (blue) substantially outperforms ipSAE (gray) for AUROC. **(D)** Pseudo-structure complexes of two *de novo* VHHs (red) confirmed with AlphaSeq as strong, specific binders to antigens (light blue) CRLF2 and ORM2.

Our classifier operates on Boltz-2’s pairformer representations (**s, z**), PAE matrix, and global confidence scores. We define the Ab–Ag interface using a 10 Å C*α*–C*α* threshold and extract single-residue embeddings for paratope and epitope positions, plus cross-interface pairwise embeddings in both directions with PAE-derived features. After MLP projection to a shared space, pairwise representations are processed by an axial transformer with factorized row/column attention, avoiding the cost of full triangular attention. Both cross-interface directions (**z**_para→epi_, **z**_epi→para_) are processed symmetrically and pooled. Aggregated features are broadcast to residue positions, concatenated with single-residue embeddings, and passed through a lightweight transformer encoder for cross-residue information flow. The output is mean-pooled, concatenated with global confidence metrics, and fed to an MLP head predicting binding versus non-binding. Full architectural details are provided in Supplementary Material E.3.

#### 2.5.2 ABACUS DND Enrichment Analysis

We compare ABACUS to ipSAE, a confidence metric previously shown to be an effective filter in DND experiments [41], using SEPIA as an in-distribution test as well as the *de novo* VHH dataset described in Section 2.3.2 (4,930 candidate VHHs against 22 antigens; 37 confirmed hits across 8 antigens) as a strict hold out test. As confidence metrics are the standard filtering paradigm for protein design, we believe this approach is representative of filtering strategies used in most modern DND pipelines.

To determine practical utility for high-throughput screening applications, we focused on enrichment metrics that quantify model performance when selecting top-ranked candidates: AUROC, AUPRC, and Enrichment@k, defined as the fold-enrichment over random selection: Enrichment@k = (Precision@k)*/*P, where P represents the baseline precision, the overall fraction of binders in the dataset. Supplementary Section F.5 shows that ABACUS excels at the in-distribution task of classifying VHH–SEP pseudo-structures as binder versus non-binder, outperforming ipSAE across all enrichment metrics.

To evaluate our ability to enrich hits in aggregate as well as for specific targets of interest, we compute metrics both globally and separately for each antigen in the *de novo* VHH dataset. Figure 5 shows global enrichment values when using either ABACUS or ipSAE to rank *in silico* designs, demonstrating approximately a 4-fold enrichment for ABACUS for *in vitro* validated hits on the top 10% of designs. Similarly, ABACUS exhibits improvement over ipSAE for AUROC, both globally and per antigen. Furthermore, we observe that the highest quantiles of ipSAE scores seem to be negatively correlated with binding (i.e. for its top quantiles, ipSAE yields worse-than-random enrichment), indicative of overconfidence [38]. Supplementary Section F.6 additionally demonstrates that training ABACUS with SEPIA data consistently outperforms training only on the PDB-derived decoy dataset. These results show that ABACUS, trained on SEPIA pseudo-structures and PDB-derived decoys, can substantially improve DND hit rates by scaling *in silico* design breadth with more effective candidate ranking.

## 3 Discussion

The central aim of this study is to address a fundamental bottleneck in *de novo* antibody design: the scarcity, bias, and slow growth of structural training data. Current structure-based models rely heavily on experimental co-structures which reflect historical experimental priorities, contain only successful binders, and expand incrementally through labor-intensive protein biochemistry and structural biology. SEPIA introduces a new paradigm: instead of passively consuming structural data, we actively generate it.

Our contributions here are threefold. First, we show that a large corpus of synthetic structural epitopes can be rapidly generated *in silico* with structure-guided DND and validated *in vitro* to produce joint sequence–structure–affinity datasets at scale. Second, we establish that the pseudo-structures of VHH–SEP complexes yielding strong, specific binding in AlphaSeq constitute sufficiently accurate structural data points to effectively train ML models for DND-relevant tasks. They are also comparable in information content and interoperable with experimentally determined co-structures for such purposes. Finally, we demonstrate that models trained with SEPIA data outperform standard confidence metrics in prioritizing successful designs within modern DND pipelines

SEPIA currently comprises approximately one thousand novel binding VHH–SEP pairs, along with many thousands of VHH point mutants, each associated with a pseudo-structure and experimental binding affinity. In this proof-of-concept study, parental VHHs were drawn from known PDB structures to maximize DND success rates. However, this is not a requirement; synthetic epitopes can just as easily be designed against other antibody formats (e.g., scFv), or against antibodies without known structures using structure prediction models. As DND success rates improve, prospective antibody design will itself become a scalable source of validated pseudo-structures. In this regime, iterative cycles of design and validation will generate increasingly informative data, creating a virtuous cycle in which improved models drive higher hit rates and enable further data generation.

In addition to simply expanding the volume of structural data available for ML model training, SEPIA data provide several distinct advantages compared to experimental co-structures already present in the PDB. One defining feature of SEPIA is the use of synthetic epitopes to explore structural space deliberately rather than incidentally. Unlike natural antigens, SEPs can be engineered to probe underrepresented or structurally novel binding modes, systematically expanding antibody-accessible epitope space beyond what is observed in the PDB. Our analyses show that SEPIA complexes occupy regions of monomer and complex structural space largely distinct from existing databases (Supplementary Material C.3; Figure S14). We attribute this finding to SEPs presenting “epitope fields” that resemble those of the natural antigens recognized by the corresponding VHHs, despite lacking overall structural similarity to these natural targets. As SEPIA expands across additional parent VHHs and paratopes, it will generate structural information progressively further from historical biology, which is precisely the regime where generalizable ML models must operate. Beyond just binary binding outcomes, SEPIA facilitates the measurement of VHH–SEP local affinity landscapes through systematic mutagenesis. These data help resolve finer patterns of antibody–antigen binding, allowing ML models to move beyond hit–miss classification toward quantitative ranking of variants, which is critical for DND campaigns in follow-up zero-shot affinity optimization. Finally, the huge number of negative-labeled designs produced as part of this work constitute a large dataset of proposed co-structures that appear high-confidence according to today’s best DND design and classification models but that do not represent true binding events. The inclusion of tens of thousands of these “hard negatives” provides a valuable source of training data that are completely absent from databases of experimental co-structures which by definition include only positive examples.

Despite its advantages, we acknowledge our approach carries several limitations. First, pseudo-structures are supported by functional binding assays rather than experimental co-crystal structures, and while strong specific binding suggests the accuracy of the designed co-structure, mutagenic characterization and functional validation alone do not guarantee atomically accurate complexes. In addition, our yeast-display system may not fully capture all biophysical contexts relevant to therapeutic development and can introduce false negatives due to proteins that may express poorly or lack mammalian post-translational modifications. SEPIA should therefore be viewed as a scalable source of structurally informed training data that augments, rather than replaces, high-resolution structural biology.

Broadly, SEPIA reflects a shift in how structural training data are generated. Traditional databases encode what nature and crystallography have made available. In contrast, pseudo-structure generation at scale allows structural datasets to be shaped by modeling needs and design objectives rather than historical serendipity. As ML performance increasingly depends on data diversity, balance, and task alignment, the ability to prospectively generate labeled co-structure data may become as central as advances in model architecture. We anticipate that large-scale, experimentally validated pseudo-structure generation—coupling *in silico* design with high-throughput *in vitro* validation—will play a defining role in the next generation of structural ML systems for antibody engineering.

## Acknowledgments

The authors acknowledge the importance of open-source software, models, and data that have been used extensively in this paper, including but not limited to RFDiffusion, ProteinMPNN, Boltz-2, numpy, PyTorch, scikit-learn, mmseqs2, FoldSeek, PDB, AFDB, SAbDab, and UniProt.

