## Supplementary Material for "The Synthetic Epitope Atlas: High-Throughput Design and Validation of *De Novo* Antibody-Antigen Complexes"

**G References** **35**

### A *In vitro* Assays

#### A.1 AlphaSeq Assay Details

To generate large-scale affinity data, AlphaSeq experiments were performed as previously described. [1, 2] Briefly, plasmids encoding yeast surface display cassettes were constructed and linearized for integration into the yeast genome. To generate VHH libraries and SEP libraries, a multiplexed gene fragment pool or oligonucleotide pool ordered from Twist Bioscience (South San Francisco, CA) was PCR amplified and inserted into pSYNAG vectors using Gibson assembly. The resulting assembled plasmids were linearized by PCR amplification to generate fragments for transformation. Parental MATa and MATalpha AlphaSeq yeast strains were co-transformed by electroporation with a mixture of the linearized DNA library fragments and a library of random barcode fragments to link each library member with multiple barcodes. To map DNA barcodes and library sequences, fragments containing the genes and associated DNA barcodes were PCR amplified from the yeast libraries and sequenced using a GridION sequencer from Oxford Nanopore Technologies (Oxford, UK). Library-on-library AlphaSeq assays were performed for each campaign by combining MATa and MATalpha libraries in YPD media with a low concentration of Tween 20 and incubating for 16 h. Control yeast strains with known interaction affinities were included as standard controls. Following induced recombination to link MATa and MATalpha barcodes on a single chromosome, the barcode locus was PCR amplified and sequenced using an Illumina NextSeq 500. Sequencing data were analyzed to identify MATa and MATalpha barcode pairs, normalized based on haploid frequencies, and assigned estimated affinities using a linear regression derived from the control strains.

### B SEPIA Experiments

This section describes the SEPIA design pipeline and experimental results across three rounds. Round 1 targeted 48 VHHs as an initial proof of concept. Round 2 expanded to 180 VHHs with pipeline modifications. Round 3 performed targeted mutational scans to validate predicted binding interfaces. Pipeline details are provided first, followed by round-specific experimental details.

#### B.1 SEPIA Design Pipeline Details

The SEPIA design pipeline generates predicted VHH–synthetic epitope protein (SEP) complexes through four sequential stages: (i) SEP backbone generation in complex with a VHH, (ii) SEP inverse folding, (iii) VHH–SEP co-structure prediction and filtering, and (iv) diversity-aware selection.

##### B.1.1 Backbone Generation with RFDiffusion

SEP backbones were generated using RFDiffusion [3], conditioned on a fixed input VHH structure. VHH structures in this work were chosen from solved crystal structures. Hotspot residues corresponding to VHH CDR positions were specified to guide interface formation during diffusion toward the paratope. SEP lengths were constrained to 50–84 amino acids. Multiple RFDiffusion configurations were evaluated during pilot testing, including standard and beta-strand–biased checkpoints, varied amounts of noise during diffusion, and both RFD-native and custom potentials (such as binder radius of gyration or a custom in-house potential that steers RFDiffusion toward structures represented with Foldseek 3Di encodings) to encourage backbone structural diversity (Section B.2).

Generated backbones were required to form at least three inter-chain contacts with the VHH. To reduce redundancy and increase diversity, backbones were clustered using Foldseek [4] at a TMScore cutoff of 0.8 prior to progressing to inverse folding, with clustering thresholds chosen based on pilot analyses of structural diversity and *in silico* pass rate (Section B.2).

##### B.1.2 Inverse Folding with ProteinMPNN

Each retained RFDiffusion backbone was inverse-folded using ProteinMPNN [5], conditioned on the VHH–SEP complex structure, to generate multiple candidate SEP sequences. ProteinMPNN temperature and sampling depth were tuned via pilot runs (Section B.2) to balance sequence diversity and downstream structural diversity and fidelity. Sequences flagged as UniRef outliers were removed prior to structure prediction (Section E.1).

##### B.1.3 Structure Prediction and Filtering with Boltz

Candidate SEP–VHH complexes were evaluated using Boltz structure prediction [6] without MSAs, with the RFDiffusion output provided as a template structure. In Round 1, Boltz-1 was used for both SEP monomer and VHH–SEP complex predictions. Designs were filtered based on TM-score between the predicted complex and the intended RFDiffusion complex, and on Boltz's aggregate confidence metrics. Unless otherwise specified, designs were required to satisfy TM-score > 0.8 and complex confidence > 0.8 to pass *in silico* filtering.

The results of SEPIA round 1 demonstrated that complex-level predictions were more informative than monomer predictions for identifying successful binders. Accordingly, Round 2 used Boltz-2 for complex prediction only, templated on the RFDiffusion output, and generated three independent complex predictions per design. Designs were retained if at least one prediction met the TM-score and confidence thresholds.

**Templating strategy.** Round 2 used Boltz-2's `force_template` option with a 3 Å distance cutoff, which constrains predicted atom positions to remain within 3 Å of the supplied template coordinates; we refer to this as *hard templating* hereafter. After SEPIA design testing was complete, subsequent analysis on SEPIA and related datasets [7] compared three Boltz-2 templating strategies: (i) no template (sequence only), (ii) soft templating (template provided as a structural reference without positional constraints), and (iii) hard templating as used in Round 2. Soft templating consistently outperformed both hard templating and the no-template baseline in discriminating true binders from non-binders, achieving higher true-positive rates across VHH targets. Hard templating, by constraining predictions close to the input template, appears to inflate confidence metrics for both binders and non-binders, reducing the discriminatory power of structure prediction–based filtering. Because this analysis was completed after Round 2 design selection, the Round 2 filtering pipeline used hard templating throughout; Round 3 was unaffected as it used Round 2 VHH-SEP hit pairs.

##### B.2 Pilot Testing and Pipeline Optimization

Prior to large-scale production, we conducted small-scale *in silico* pilot experiments to optimize pipeline parameters. These experiments varied RFDiffusion noise levels, backbone sampling depth, ProteinMPNN sequence sampling, and filtering strategies, and evaluated outcomes using (i) *in silico* pass rate under TM-score and confidence thresholds and (ii) structural diversity assessed via Foldseek clustering. Based on these pilots, Round 2 prioritized increased backbone diversity (more RFDiffusion backbones with fewer sequences per backbone, reduced from 10 to 3), removed monomer-level filtering, and introduced a custom in-house 3Di encoding-based RFDiffusion checkpoint alongside the standard and beta-strand–biased checkpoints. In Round 2 production, designs were split approximately equally across standard, beta-strand–biased, and 3Di encoding-based checkpoints (each contributing roughly one-third of generated backbones).

The final production configuration (Round 2 onward) used the following parameters. Total design sampling depth through the pipeline was set to approximately 20× the intended number of SEPs per VHH, ensuring sufficient candidates after filtering. RFDiffusion employed standard and beta-strand–biased checkpoints, noise scale and noise scale frame both set to 0.2, and the binder radius of gyration potential from the RFDiffusion repository with default settings (weight 1, guide scale 1, guide decay constant). A subset of RFDiffusion designs additionally applied our in-house custom potential to guide diffusion toward a random selection from a library of FoldSeek 3Di encodings [4] of small natural proteins, using a stochastic policy gradient estimation for the gradient with respect to backbone atomic coordinates. Each backbone was inverse-folded with ProteinMPNN at a sampling temperature of 0.1, generating 3 candidate sequences per backbone with cysteine excluded from proposals. Each candidate was evaluated with 3 templated Boltz-2 complex predictions. Interface pseudo-structure analysis used the default `ipSAE_max` variant of the interaction prediction Score from Aligned Errors (ipSAE; [8]) — the maximum of the two asymmetric align-on-$A$/score-$B$ values — with default settings (PAE cutoff 10, ipSAE distance cutoff 10). Round 1 differed in that it used Boltz-1 for both monomer and complex predictions (Section B.4.1).

**Binding classifier for design selection.** To prioritize SEP candidates for experimental testing, we trained an XGBoost classifier to predict binding probability from structure prediction and sequence features. The model was trained on five diverse AlphaSeq datasets (∼87,000 PPIs) using 5-fold cross-validation with splits on unique target proteins. Input features comprised Boltz-2 complex confidence metrics (pTM, iPTM, pLDDT, iPLDDT, PDE, iPDE, confidence score), interface-derived metrics (ipSAE, pDockQ, pDockQ2, LIS, and related scores), and ProtParam physicochemical properties of both binding partners. The classifier was applied as a soft ranking filter as part of Round 2 SEP selection (Section B.4.2), not as a hard pass/fail threshold.

##### B.3 Definition of a Hit

We define a VHH–SEP interaction as a hit based on absolute binding strength and specificity relative to off-target interactions. Because AlphaSeq measures all pairwise interactions between the VHH and SEP libraries, off-target binding can be directly quantified and incorporated into hit-calling. AlphaSeq provides calibrated binding affinity estimates ($K_d$ $\log_{10}$ nM) for each pairwise VHH–SEP interaction, which we use to identify strong, specific binders via the criteria below.

**Off-target statistics.** For each SEP, we compute robust statistics over its off-target interactions (interactions with VHHs that the SEP was *not* designed to bind), using only interactions with sufficient barcode counts:

- $a^{\text{off}}_{\text{min}}$: minimum off-target predicted affinity (strongest off-target binder)
- $\tilde{a}^{\text{off}}$: median off-target predicted affinity
- $\hat{\sigma}^{\text{off}} = 1.4826 \times \text{MAD}(a^{\text{off}})$: robust standard deviation estimate, where MAD is the median absolute deviation and the scaling factor ensures consistency with the normal distribution
- $a^{\text{off}}_{3\sigma} = \tilde{a}^{\text{off}} - 3\hat{\sigma}^{\text{off}}$: three-sigma threshold below the off-target distribution

**Hit criteria.** A VHH–SEP pair is classified as a hit if all four criteria in Table S1 are satisfied.

**Table S1:** Criteria for classifying a VHH–SEP binding hit.

| # | Criterion | Definition |
|---|---|---|
| 1 | Data quality | Sufficient barcode counts for reliable affinity estimation |
| 2 | Affinity | AlphaSeq affinity $< 1\ \mu$M |
| 3 | On-target | SEP was explicitly designed against this VHH or a mutant thereof |
| 4 | 3-sigma significance | On-target affinity is more than three robust standard deviations below the off-target median: $a_{\text{on}} < a^{\text{off}}_{3\sigma}$ |

##### B.4 Details of SEPIA Rounds

###### B.4.1 SEPIA Round 1 Details

We designed SEPs against 48 VHHs selected from SAbDab-nano, [9, 10] chosen to maximize diversity in CDR3 length and composition (Table S2). The pipeline generated ∼600,000 candidate SEPs (∼12,500 per VHH). SEPs were selected for AlphaSeq testing by applying a global filter (TM-score $> 0.8$ and Boltz-1 complex confidence $> 0.8$) agnostic to VHH identity, yielding 25,448 SEPs (∼4% *in silico* pass rate). As this was the first round of SEPIA pipeline testing, we intentionally oversampled SEPs for a subset of VHHs to ensure sufficient hits for downstream analysis in case hit rates were low. After *in silico* filtering, per-VHH SEP counts ranged from 48 to 3,158 (median 356; Figure S1).

AlphaSeq screening of the 25,448 SEPs against the 48 VHHs produced ∼1.2 million pairwise protein–protein interaction (PPI) measurements. We identified 928 VHH–SEP interactions ("hits") satisfying the criteria defined in Section B.3, corresponding to an *in vitro* success rate of 3.6% of tested SEPs. Of the 48 VHHs tested, 34 (70.8%) yielded at least

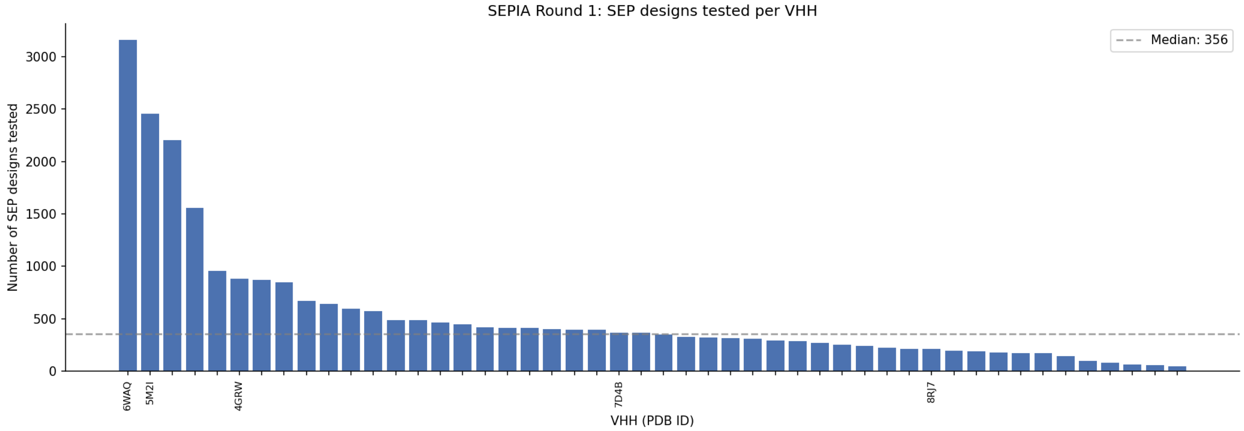

**Figure S1:** Number of SEPs tested per VHH in SEPIA Round 1. SEPs were selected using a global structure-based filter (TM-score $> 0.8$, Boltz-1 complex confidence $> 0.8$) applied uniformly across all VHHs, resulting in variable per-VHH counts reflecting differences in *in silico* pass rates. Dashed line indicates median (356 SEPs).

**Table S2:** PDB IDs of 48 SEPIA Round 1 VHHs.

| |
|---|
| 1KXQ, 1MEL, 2VYR, 4EIZ, 4GRW, 4KML, 4N1H, 4NC1, 4ORZ, 4QO1, 4S10, 4Y7M, 5C2U, 5E5M, 5G5X, 5IMM, 5LHN, 5M2I, 5M2M, 5N88, 5O02, 5Y7Z, 6EY0, 6FE4, 6GWP, 6OQ5, 6U54, 6WAQ, 6XZU, 7B5G, 7D4B, 7E53, 7KJI, 7N0R, 7NDF, 7VFB, 7XLD, 8DQU, 8EW6, 8F6V, 8H63, 8J5J, 8K3A, 8PE1, 8PIH, 8Q7S, 8RJ7, 8YJ5 |

one validated SEP hit. Success rates were highly variable across VHHs: some VHHs had dozens of validated binders while others engaged only one or two hits.

##### B.4.2 SEPIA Round 2 Details

SEPIA Round 2 expanded the target set to 180 parental VHHs from SAbDab-nano (Table S3), comprising 38 VHHs previously tested in Round 1 and 142 new VHHs introduced to broaden paratope diversity. These VHHs and their corresponding point mutants were drawn from a prior large-scale AlphaSeq survey of SAbDab-nano VHH–antigen interactions [11], which included targeted CDR mutational scans for the same 180 parental VHHs.

**VHH mutant generation.** In addition to parental VHHs, we included 11,425 computationally designed VHH point mutants to assess SEP binding to VHH sequence variants. Mutants were generated using a multi-stage filtering pipeline:

*Single-site mutagenesis (SSM).* For each parental VHH, we generated all possible single-amino-acid substitutions at every position, resulting in $\sim$600,000 candidate sequences across all parental VHHs.

*ESM probability filtering.* We filtered candidates using ESM-2 pseudo-log-likelihood scores, retaining only mutations where the mutant probability exceeded a threshold of $\log(p_{\text{parental}}) - 2$, to enrich for tolerated mutations, which reduced the candidate pool to $\sim$225,000 sequences.

*PSSM filtering.* We constructed a position-specific scoring matrix (PSSM) from all VHH sequences in SAbDab-nano using ANARCI numbering. Candidates with substitutions to amino acids occurring in fewer than 2.5% of SAbDab-nano VHHs at that position were filtered out, primarily removing mutations in framework regions unlikely to impact binding while potentially affecting expression or stability. This reduced the pool to $\sim$100,000 sequences.

*Redundancy reduction.* To maximize sequence diversity within our sequence budget, for a given VHH mutation we cap the number of VHHs we apply this to be at most 5, reducing the number of duplicate substitutions across parental VHHs.

*Control sequences.* A small number of filtered-out sequences were added back as negative controls to assess filter calibration.

The final set comprised 11,425 unique VHH mutants ($\sim$50–100 per parental), enabling assessment of how single-residue changes affect SEP binding specificity across diverse parent VHHs.

**Design scale and filtering.** The Round 2 pipeline generated $\sim$660,000 candidate SEPs (220,000 unique backbones $\times$ 3 ProteinMPNN sequences each). Candidates were prefiltered by requiring both TM-score $> 0.8$ (structural agreement between the Boltz-2 complex prediction and the designed RFDiffusion backbone) and Boltz-2 complex confidence $> 0.8$, yielding 43,324 unique SEPs corresponding to 28,027 unique backbones.

**SEP selection for testing.** 43,324 SEPs passed initial SEPIA pipeline filters used in Round 1 (TMscore and Boltz-2 complex confidence > 0.8). To reduce structural redundancy, the 43,324 prefiltered SEPs were clustered using Foldseek `multimercluster` at a TM-score threshold of 0.8, yielding 24,428 unique complex clusters; from each multi-member cluster, the design with the highest classifier binding probability was retained (Section B.2).

From this deduplicated pool, up to 80 designs per VHH were selected by descending classifier binding probability. For VHHs with fewer than 80 prefiltered designs, the allocation was supplemented with designs that did not pass prefiltering, again ranked by classifier probability. An additional 10 randomly selected unfiltered designs per VHH were included as controls to assess filter calibration. Finally, $\sim$1,000 Round 1 SEP hits were included for re-testing against Round 2 VHH point mutants to assess binding interface sensitivity. The final set comprised $\sim$22,000 SEPs, distributed across multiple AlphaSeq experiments subject to network size constraints.

**Experimental configuration.** Each SEP was tested against all VHHs in its assigned pool (both parental VHHs and point mutants thereof), generating $\sim$26 million pairwise interaction measurements.

**Table S3:** PDB IDs of 180 SEPIA Round 2 parental VHHs.

| Sub-pool | PDB ID List |
|---|---|
| 1 | 5H8O, 7O06, 2X89, 6GJU, 8QF4, 8QF5, 7DV4, 6RPJ, 6RQM, 7VKE, 5F21, 5F1O, 5F1K, 7WKI, 6Z1V, 6Z20, 4GRW, 7Z1X |
| 2 | 5IP4, 5FV2, 6RTY, 6RVC, 8AOK, 7CZD, 5JDS, 8AOM, 7EOW, 5DMJ, 6R7T, 1OP9, 7D4B, 9EN2, 8B7W, 4TVS, 7O3B, 8UO9, 7ANQ, 8OZB |
| 3 | 4DK6, 4YGA, 6FV0, 7UNZ, 3OGO, 3G9A, 8G0I, 8SFZ, 6LZ2, 7CZ0, 6XZF, 6UKT, 8C5H, 6X05, 6X04, 6X08, 8AV2, 7UST, 7USV, 7R24, 4GFT, 8E0E, 8DAM, 3STB |
| 4 | 7UBX, 4NC2, 7YZI, 8ILX, 8IM0, 8IM1, 7SAJ, 8RWF, 5DA0, 6DBF, 6U12, 6RAL, 9G5Z, 9G48, 8R4B, 7XLD |
| 5 | 6QV2, 4WEN, 8C3K, 5JA9, 5JA8, 9FVB, 9FVC, 5MP2, 4WEU, 5VXL, 5VXJ, 5VXM, 5VXK, 5OVW, 4W6X, 4W6W, 4W6Y, 3CFI |
| 6 | 7RI1, 4W2O, 4W2Q, 6U51, 6U55, 8WCG, 7X2J, 6WAQ, 6XW5, 6XW7, 8YSF, 8YSH, 8IDM, 8IEE, 5BOP, 7NFT, 2BSE |
| 7 | 5Y7Z, 5N88, 8Z8V, 4KML, 6GWP, 6ZRV, 4S10, 8K3A, 8Z8M, 5M2M, 5M2J, 8W90, 8EW6, 4QO1, 9FWW, 2VYR |
| 8 | 7E53, 8PE1, 8PE2, 8PIH, 1KXT, 1KXQ, 1MEL, 1ZVY, 1XFP, 1RJC, 6XZU, 5LHN, 5IMM, 5C2U |
| 9 | 8F6V, 8EVD, 8ELN, 8GJR, 6EY0, 4NC1, 8YJ5, 6OQ5, 5LWF, 4Y7M, 4M3K, 4N1H, 7B5G, 4EIZ, 4EIG, 3K74, 4I13, 8H63, 8H64, 5G5X |
| 10 | 6U54, 8Q7S, 8CYC, 8CYD, 8BEV, 4ORZ, 5O02, 7VFB, 7N0R, 7R98, 8DQU, 2XXM, 7N9V |

###### B.4.3 SEPIA Round 3 Details

SEPIA Round 3 was designed to validate VHH–SEP binding interfaces through systematic mutagenesis, comprising two complementary experiment types.

**SEP site saturation mutagenesis (SSM).** We selected a validated SEP design targeting the 7XLD_B parent VHH and performed a comprehensive site-saturation mutagenesis (SSM). The parental SEP was mutated at every position to all 18 alternative amino acids (excluding cysteine), plus random pair combinations, yielding 4,779 SEP variants in total. These variants were screened against the wild-type 7XLD_B VHH using AlphaSeq. Of 4,779 variants tested, 3,496 (73.2%) met our hit criteria, enabling identification of positions tolerant or intolerant to mutation. Details of the SSM analysis are provided in Section C.4.3.

**Double-sided alanine scanning.** To directly validate predicted contact interfaces, we performed systematic alanine scanning mutagenesis on both VHH and SEP binding partners. We selected 4 parent VHHs (4NC1_E, 5M2I_J, 6U54_A, 6XZU_A) and generated CDR3 alanine-scan libraries for each, yielding 57 unique VHH sequence variants (including wild types). Wild-type alanine residues were mutated to serine residues in the scans. We then generated alanine scan variants for 24 SEP hits for each of the 4 parent VHHs, yielding 7,040 unique SEP sequence variants. The resulting AlphaSeq dataset comprises 114,187 on-target pairwise measurements. We identified 35,010 hits, with 56 VHH variants (98.2%) and 5,488 SEP variants (78.0%) participating in at least one hit interaction. Details of the alanine scan analysis are provided in Section C.4.2.

#### C Analysis of SEPIA Results

This section characterizes validated wild-type VHH–SEP hits from Rounds 1 and 2 across three analyses: (i) physicochemical and sequence-level statistics of hit SEPs, (ii) epitope contact frequency analysis, and (iii) structural diversity and PDB novelty assessed via Foldseek [4] at both the monomer and complex levels.

##### C.1 SEP Statistics

We characterized the physicochemical and structural properties of all wild-type on-target SEPs across Rounds 1 and 2. After deduplication to one entry per unique SEP, the analysis covers 45,430 unique SEPs (1,161 hits, 44,269 non-hits). Properties were computed using BioPython ProtParam [12] on unique SEP sequences.

**Table S4:** Summary of SEP physicochemical properties computed via BioPython ProtParam for all unique wild-type on-target SEPs.

| **Property** | **Mean ± Std** | **Min** | **Median** | **Max** |
|---|---|---|---|---|
| Sequence length (aa) | $76.4 \pm 4.0$ | 70.0 | 76.0 | 84.0 |
| Molecular weight (Da) | $8112.2 \pm 651.4$ | 5562.0 | 8094.2 | 10384.7 |
| Isoelectric point (pI) | $5.48 \pm 1.44$ | 4.05 | 4.95 | 12.00 |
| GRAVY | $-0.24 \pm 0.59$ | $-2.38$ | $-0.21$ | 2.38 |
| Instability index | $33.36 \pm 16.07$ | $-14.25$ | 31.88 | 133.25 |
| Aromaticity | $0.03 \pm 0.02$ | 0.00 | 0.03 | 0.19 |
| Charge at pH 7 | $-3.65 \pm 3.46$ | $-23.09$ | $-3.50$ | 22.75 |

##### C.1.1 SEP Physicochemical Properties

Table S4 summarizes the physicochemical properties of all 45,430 unique wild-type on-target SEPs. Designed SEPs range from 70 to 84 amino acids in length (mean $76.4 \pm 4.0$ aa, mean MW $8{,}112 \pm 651$ Da), with mean isoelectric point $5.5 \pm 1.4$, slight hydrophilicity (mean GRAVY $= -0.2 \pm 0.6$), and predicted stability (mean instability index $= 33.4 \pm 16.1$, below the instability threshold of 40).

The distributions of six physicochemical properties are largely overlapping between hit and non-hit SEPs, indicating that hit status is not determined by bulk sequence-level properties (Figure S2).

Amino acid frequencies are highly similar between hit and non-hit SEPs, with no single residue strongly enriched or depleted in hits (Figure S3).

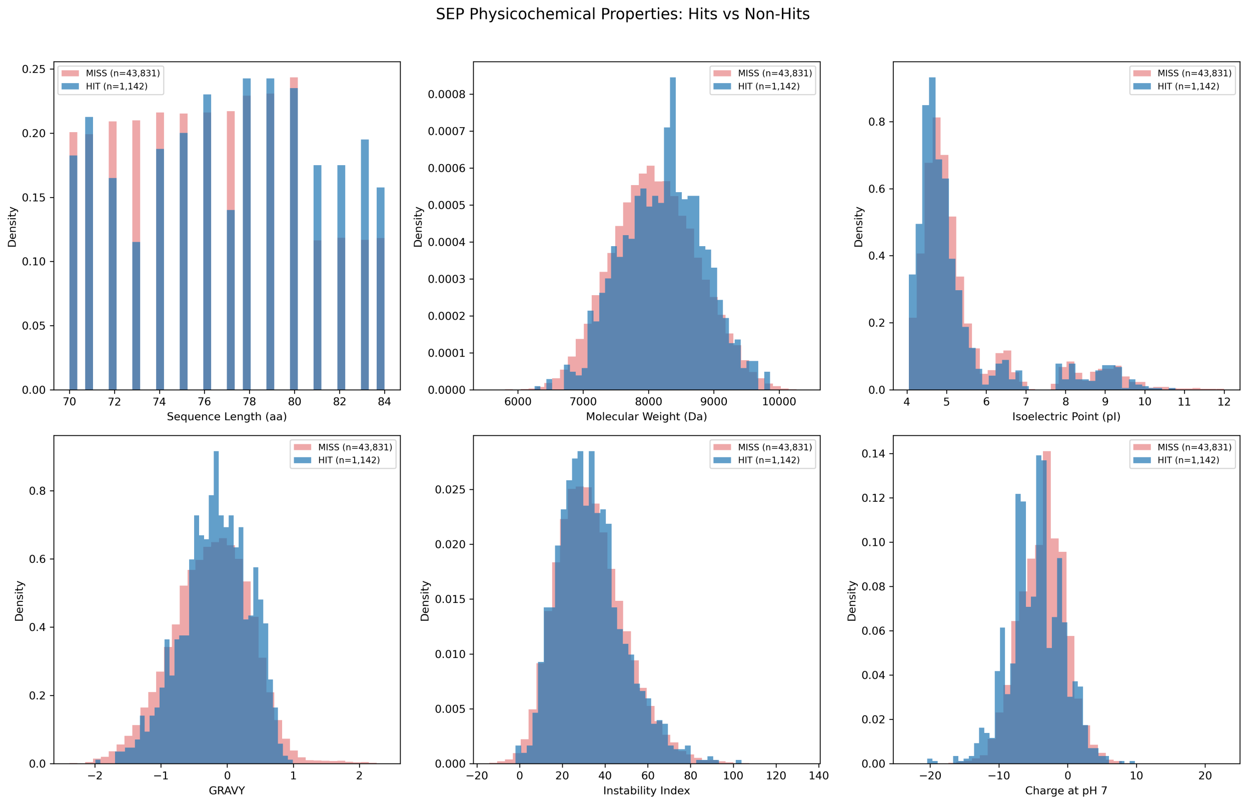

**Figure S2:** Physicochemical property distributions for hit (blue) vs non-hit (red) SEPs. Distributions are broadly overlapping across all properties.

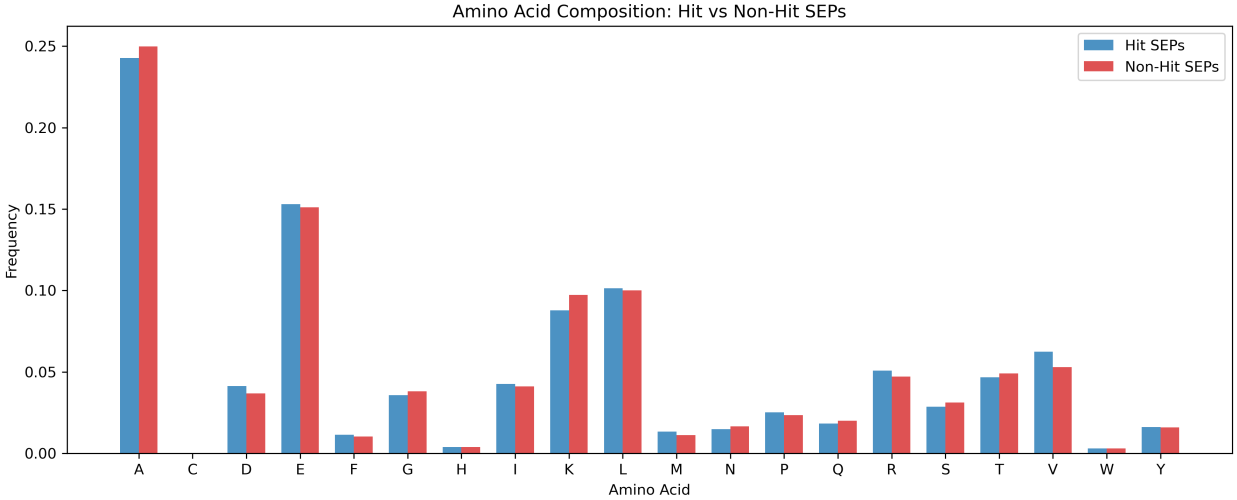

**Figure S3:** Amino acid composition comparison between hit and non-hit SEPs. Frequencies are computed per-residue across all sequences in each group.

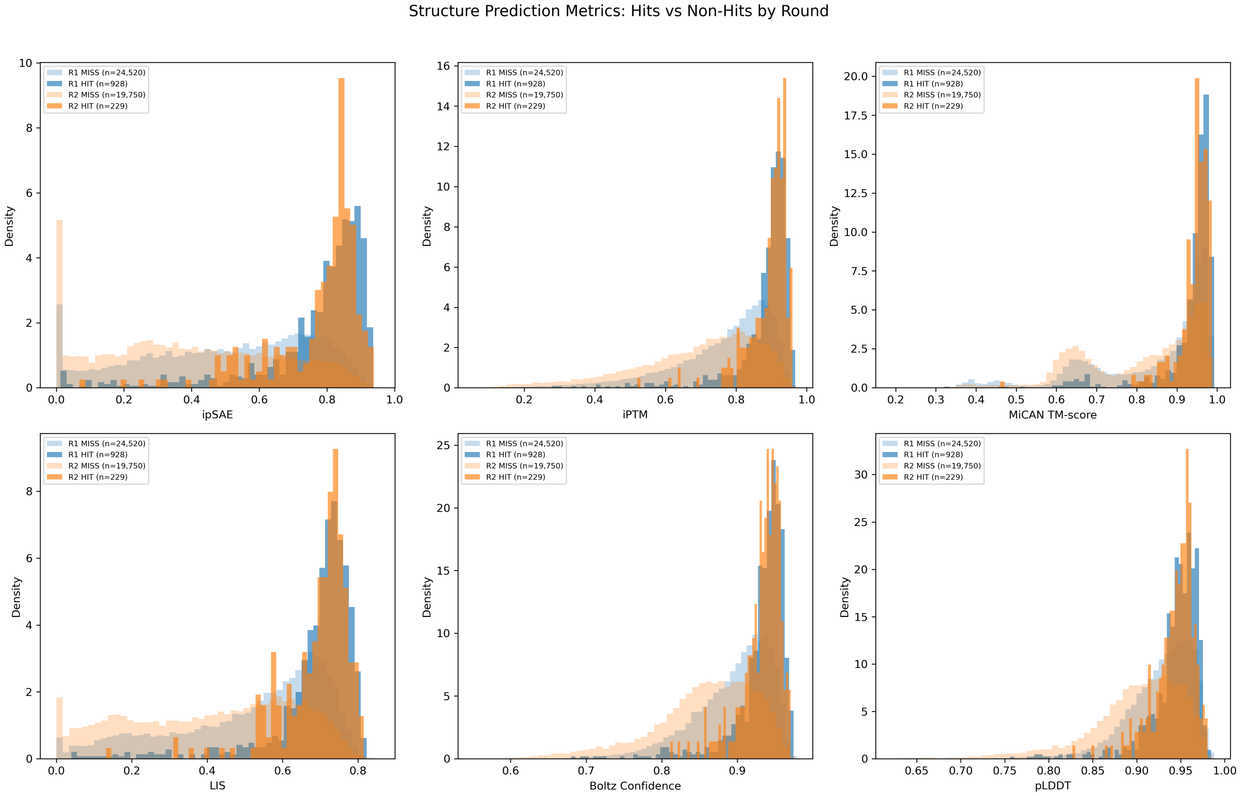

**Figure S4:** Structure prediction metric distributions for unique wild-type on-target SEPs, split by round and hit status (Round 1: 928 hits; Round 2: 233 hits, `sepia_rd2` designs only). Hits tend toward higher metric values, particularly for ipSAE and iPTM.

##### C.1.2 Structure Prediction Metric Distributions

We examined the distributions of structure prediction quality metrics for all unique wild-type on-target SEPs, stratified by round (Round 1/Round 2) and hit status. Metrics include ipSAE, iPTM, MiCAN TM-score, LIS, Boltz confidence score, and pLDDT (Figure S4).

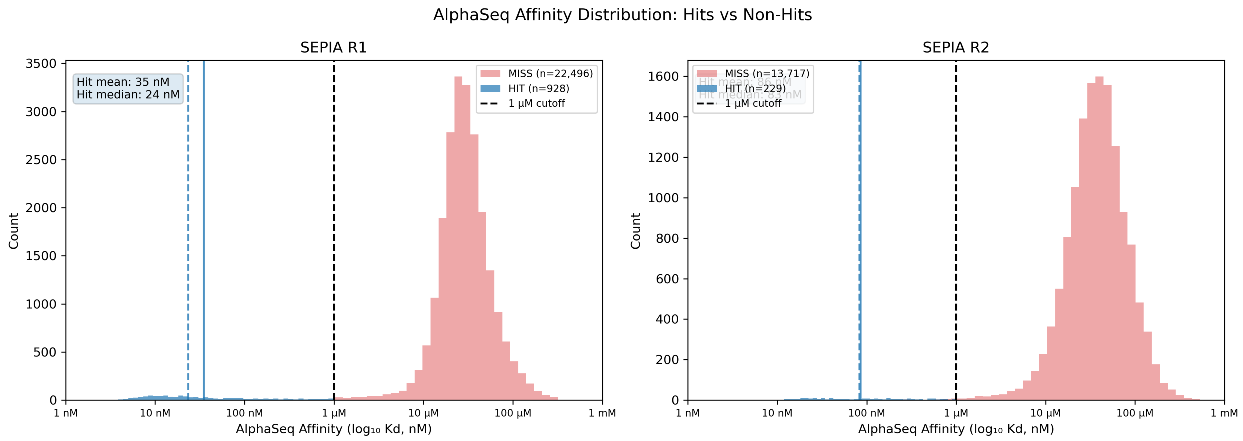

**Figure S5:** AlphaSeq affinity distributions for unique wild-type on-target SEPs in Round 1 (928 hits) and Round 2 (233 hits, `sepia_rd2` designs only). Hits (blue) are concentrated below the 1 $\mu$M threshold (dashed line).

**Note on structure prediction methodology.** As described in the main text (Section 2.1), the SEPIA design pipeline evolved between Rounds 1 and 2. In Round 1, candidate SEPs were filtered using Boltz-1 predictions (TM-score $> 0.8$, complex confidence $> 0.8$); Round 2 upgraded to Boltz-2 with RFDiffusion-backbone templates and no MSA generation. All structure prediction metrics shown in this section were recomputed uniformly using the latest pipeline configuration: Boltz-2 complex predictions, no MSAs, templated with the RFDiffusion-produced SEP–VHH complex backbone. This explains why some SEPs that passed earlier in-silico filters (based on Boltz-1 or non-templated predictions) may fall below the latest pipeline thresholds for metrics such as MiCAN TM-score or Boltz confidence.

##### C.1.3 AlphaSeq Affinity and Structure Metric Correlations

AlphaSeq affinities for unique wild-type on-target SEPs show hits concentrated at stronger binding in both Rounds 1 and 2 (Round 2 restricted to `sepia_rd2` designs; Figure S5). Across all 1,161 validated hits, the median on-target affinity is 29 nM (1.47 $\log_{10}$ nM) with an interquartile range of 13–125 nM, well below the 1 $\mu$M hit threshold. 72% of hits bind below 100 nM.

Hits cluster at strong binding affinity and high ipSAE, though many non-hits also achieve high ipSAE scores (Figure S6).

While hits are enriched at higher structure prediction metric values, no individual metric alone is sufficient to discriminate hits from non-hits (Figure S7).

#### C.2 SEPIA Naturalness Analysis

To assess whether SEPIA-designed VHH–SEP interfaces recapitulate the amino acid preferences observed at experimentally solved VHH–antigen interfaces, we computed residue-level frequency statistics at predicted binding contacts for both SEPIA Round 2 designs and the on-target SAbDab-nano structures used in the SAbDab-nano Decoy Dataset.

**Interface contact definition.** Interface contacts were identified using a symmetric nearest-neighbor procedure based on C$\alpha$–C$\alpha$ inter-chain distances. Given a predicted or experimental VHH–antigen (or VHH–SEP) complex structure, the full C$\alpha$–C$\alpha$ distance matrix $D_{ij}$ was computed between VHH residue $i$ and antigen/SEP residue $j$. Contacts were identified by scanning in both directions: for each VHH residue, the nearest antigen residue within a 10.0 Å threshold was recorded, and vice versa. The union of contacts from both scans was deduplicated by residue-index pair $(i, j)$ to avoid double-counting, yielding a set of unique paratope–epitope residue contacts per structure.

**Frequency metrics.** We analyzed two frequency metrics:

1. **Epitope contact residue frequencies:** The fraction of each amino acid among antigen/SEP residues participating in at least one interface contact, normalized to sum to 1.
2. **Paratope–epitope pair frequencies:** A $20 \times 20$ matrix $P_{ab}$ recording the joint frequency of paratope amino acid $a$ contacting epitope amino acid $b$, normalized so that $\sum_{a,b} P_{ab} = 1$.

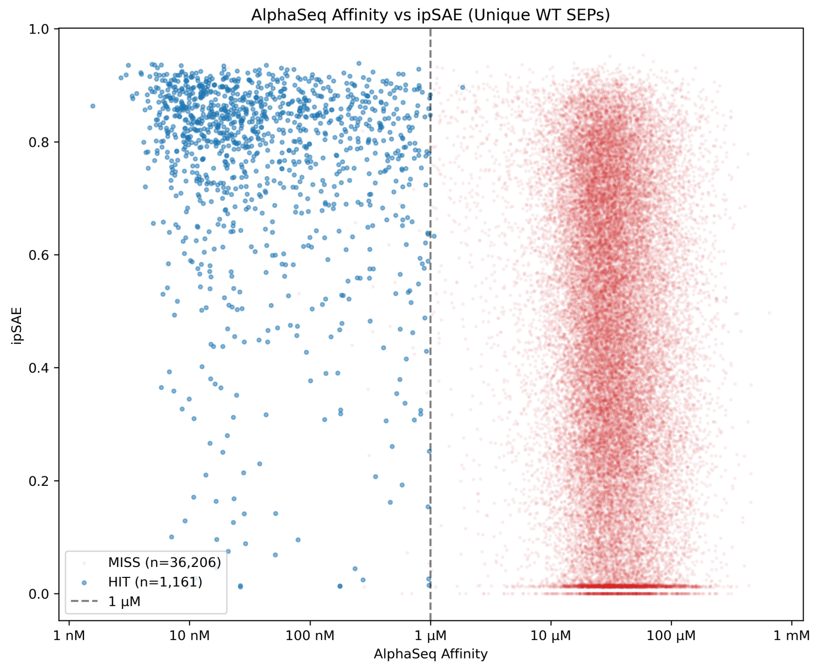

**Figure S6:** AlphaSeq affinity versus ipSAE for all unique wild-type on-target SEPs (1,161 hits, 44,269 non-hits). Hits (blue) cluster at strong binding affinity and high ipSAE.

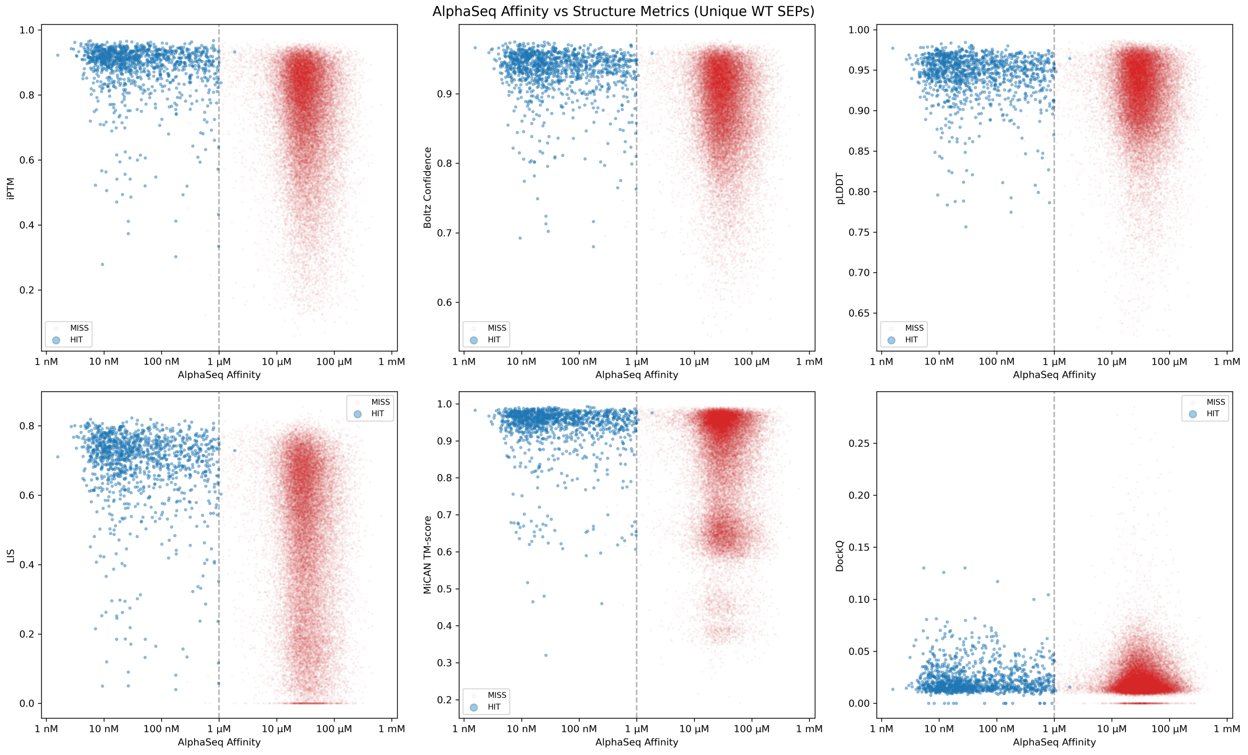

**Figure S7:** AlphaSeq affinity versus structure prediction metrics for unique wild-type on-target SEPs (1,161 hits, 44,269 non-hits). Hits (blue) are enriched at higher metric values but substantial overlap exists with non-hits.

**Epitope Frequency Distributions** We analyzed epitope contact amino acid frequencies stratified by RFDiffusion checkpoint (standard, beta, and 3Di) and compared them to the SAbDab-nano reference (Figure S8A). Briefly mentioned above in Section B.2, the 3Di SEP designs were made with the RFDiffusion standard checkpoint but using a custom potential for diffusion guidance to steer toward selected 3Di encodings of natural proteins. SEP epitope contacts showed a pronounced alanine bias, particularly in standard and 3Di designs. Cysteines were excluded by design.

For each VHH, epitope contact amino acid counts from all predicted complex structures were pooled into a single 20-element frequency vector and normalized to a probability distribution $p$. The SAbDab-nano reference was similarly represented as a single aggregate distribution $q$ over the 20 amino acids. We then computed the KL divergence $D_{\mathrm{KL}}(p\|q)$ (with $\epsilon$-smoothing, $\epsilon = 10^{-10}$) for each target VHH, yielding one divergence value per VHH. The resulting collection of per-VHH KL divergence values was visualized as distributions stratified by RFDiffusion backbone-generation strategy in Figure S8B. Here we see that the beta checkpoint produced consistently more natural designs than the alternatives.

At the pair level, paratope–epitope frequency matrices were averaged across all target VHHs and compared between SEPIA and SAbDab-nano. In addition to comparing raw pairwise distributions we also compute a difference heatmap $P_{ab}^{\mathrm{SEPIA}} - P_{ab}^{\mathrm{SDN}}$, displayed with a diverging color scale centered at zero and accompanied by marginal bar charts showing the directional difference in epitope and paratope amino acid usage. In Figure S9 we observe that within our observed epitope alanine bias, alanine-serine and alanine-glycine are particularly overrepresented epitope-paratope combinations in our SEPIA Round 2 dataset.

##### C.3 VHH–SEP Structure Analysis

We used Foldseek [4] to characterize the structural diversity of validated VHH–SEP hits and assess their novelty relative to known structures in the Protein Data Bank (PDB). Analysis was performed at two levels: (i) SEP monomer structures (chain B only) and (ii) VHH–SEP complex structures. All 1,161 validated wild-type hit structures from Rounds 1 and 2 were included in both analyses.

###### C.3.1 SEP Monomer Structural Clustering

**Methods.** SEP monomer structures (chain B) were extracted from the 1,161 predicted VHH–SEP complexes and clustered using Foldseek `easy-cluster` with a TM-score threshold of 0.8 and sensitivity 9.5. This threshold groups structures sharing the same overall fold topology.

**Results.** Clustering at TM-score $\geq 0.8$ yielded **278 unique SEP fold clusters** from 1,161 input structures (76.1% redundancy; average cluster size 4.2). The cluster size distribution is highly skewed: many clusters contain a single structure, while the largest clusters contain tens of members (Figure S10). This indicates that while the SEPIA design pipeline produces a moderate diversity of SEP fold topologies, many SEPs share similar backbone geometries, likely reflecting preferences of the RFDiffusion backbone generation step for certain helical bundle and mixed $\alpha/\beta$ architectures.

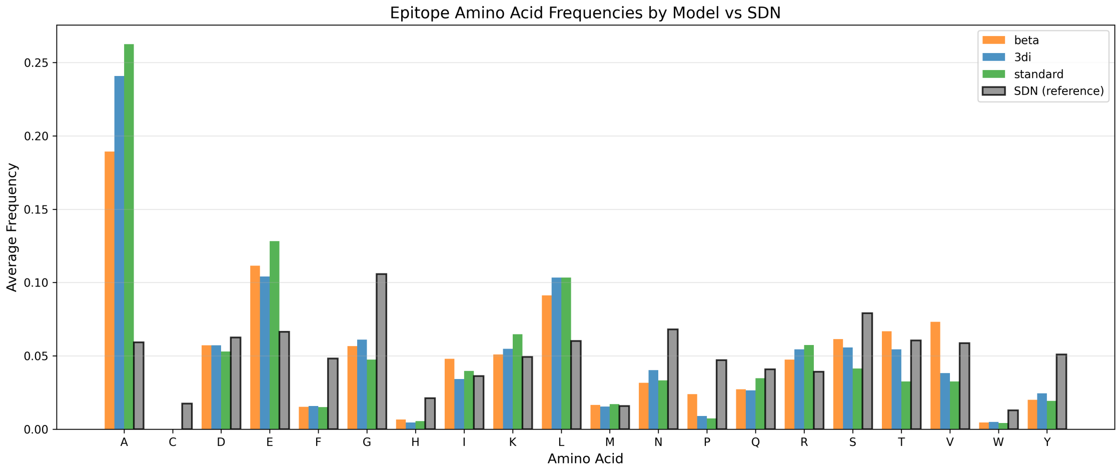

**(A)**

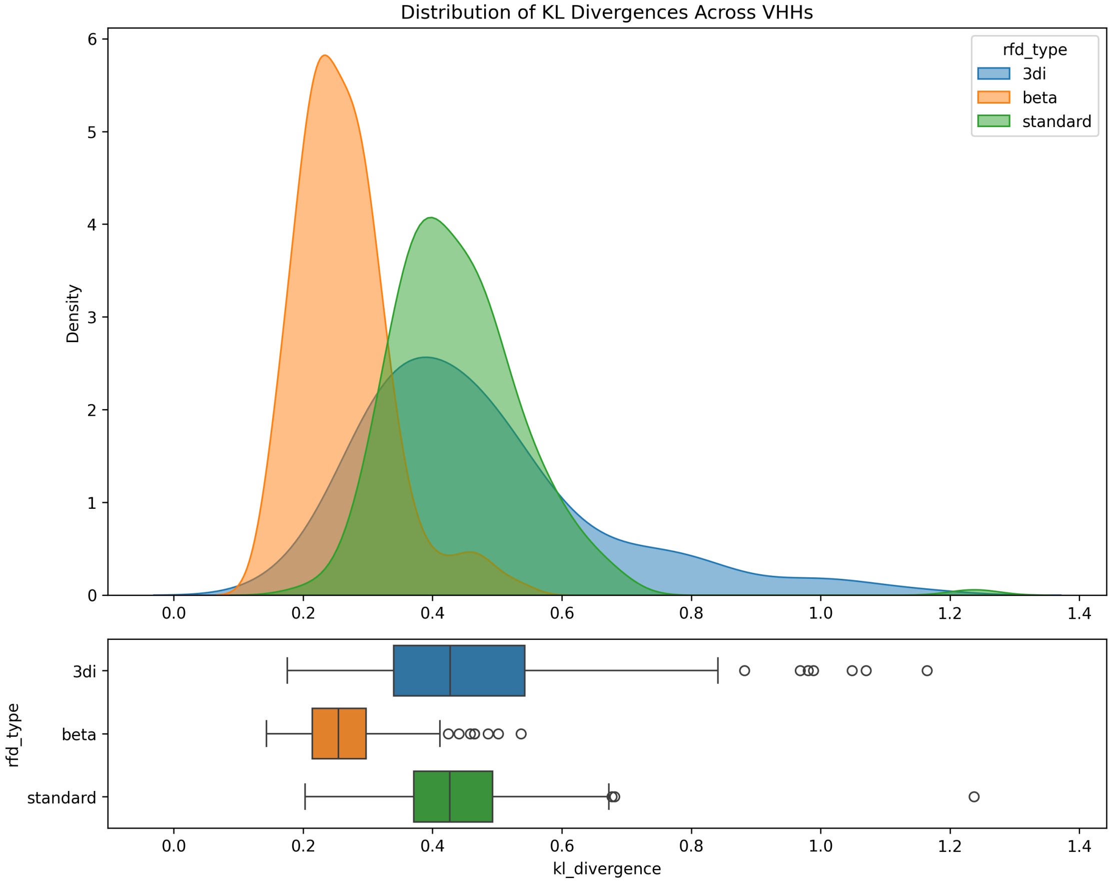

**(B)**

**Figure S8: Epitope contact amino acid distributions of designed VHHs. (A)** Amino acid frequency distributions of epitope contacts for SEPs, stratified by RFDiffusion backbone-generation strategy (standard, beta, and 3di), compared to the SAbDab-nano reference. A pronounced alanine bias particularly is visible in the standard and 3di designs, while the beta checkpoint ameliorates this to an extent. Cysteines were excluded by design. **(B)** Per-VHH KL divergence ($D_{\mathrm{KL}}(p\|q)$) of epitope contact distributions relative to the SAbDab-nano reference, stratified by RFDiffusion strategy. The beta checkpoint yields consistently lower divergence values, indicating more naturalistic epitope contact profiles.

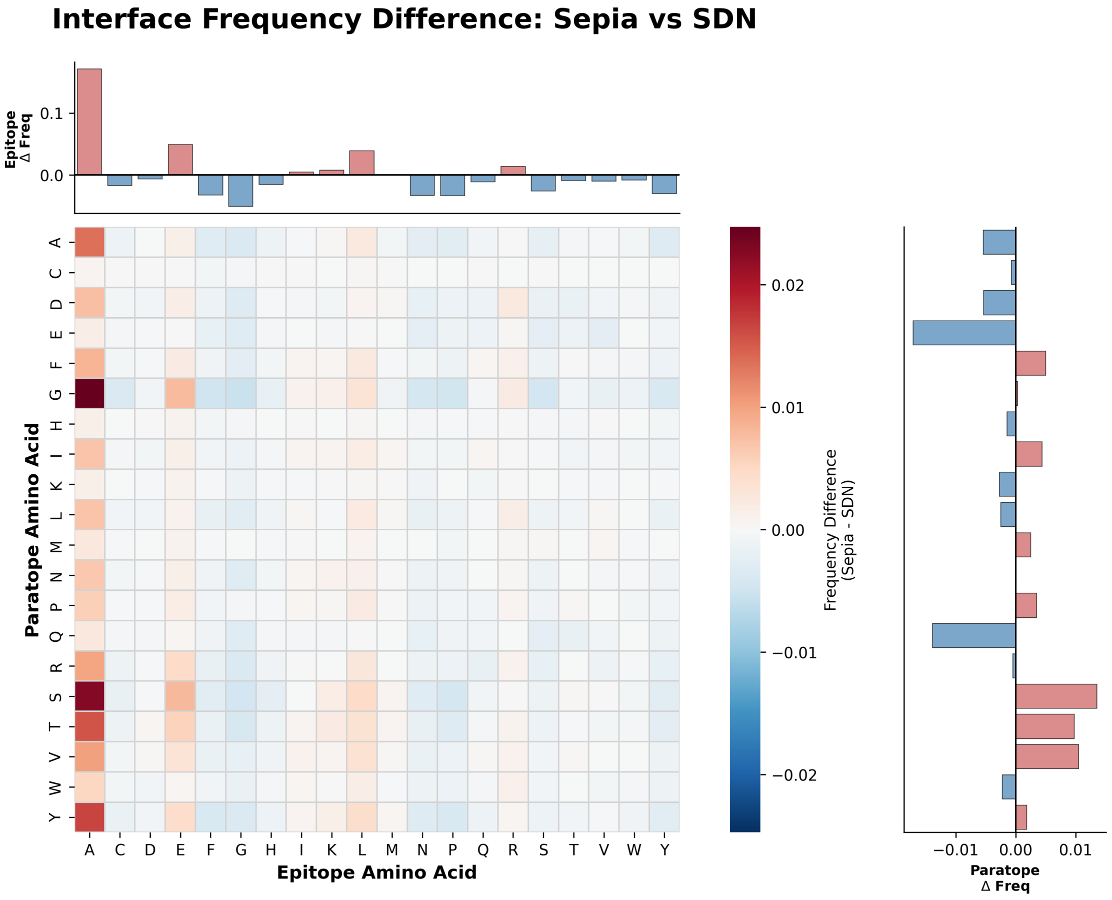

**Figure S9: Pairwise paratope–epitope frequency difference between SEPIA and SAbDab-nano.** Difference heatmap $P_{ab}^{\mathrm{SEPIA}} - P_{ab}^{\mathrm{SDN}}$ of paratope–epitope amino acid pair frequencies, averaged across all target VHHs. Red cells indicate overrepresentation in SEPIA designs; blue cells indicate underrepresentation relative to the SAbDab-nano reference. Marginal bar charts show the net directional difference in epitope (top) and paratope (right) amino acid usage.

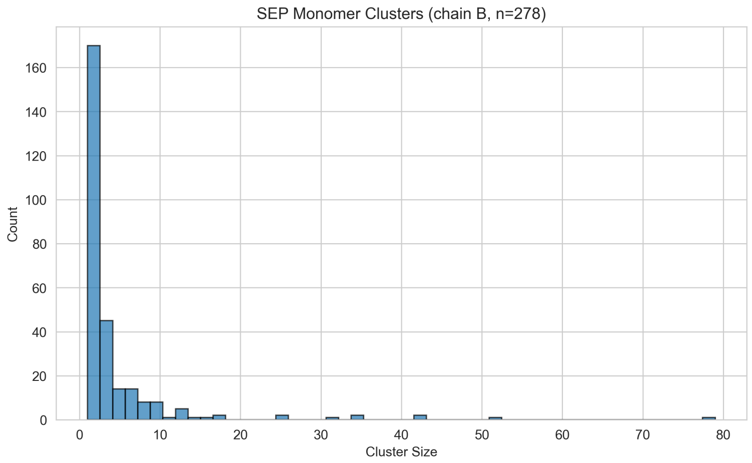

**Figure S10: SEP monomer structural clustering.** Distribution of cluster sizes for 1,161 SEP monomers clustered at TM-score $\geq 0.8$.

**Table S5: Top 5 SEP monomer PDB matches** ranked by query-normalized TM-score (`qtmscore`).

| VHH Target | PDB Hit | qTM | qCov | E-value |
|---|---|---|---|---|
| 6U54 | 3U8V_A | 0.846 | 0.973 | 0.057 |
| 6U54 | 3U8V_A | 0.835 | 0.960 | 0.036 |
| 6U54 | 3U8V_A | 0.830 | 0.960 | 0.088 |
| 5M2J | 6WI5_B | 0.819 | 0.960 | 0.003 |
| 4NC1 | 1UJW_B | 0.810 | 0.963 | 0.098 |

##### C.3.2 SEP Monomer PDB Similarity Search

**Methods.** Each of the 1,161 SEP monomer structures was searched against PDB100 using Foldseek `easy-search` with default sensitivity settings. The query-normalized TM-score (`qtmscore`) was used as the primary similarity metric.

**Results.** Only **205 of 1,161 SEP monomers** (17.7%) returned any PDB hit. Among those with hits, the median best TM-score was 0.568 and the maximum was 0.846. Twenty-six SEP monomers (12.7% of those with hits) achieved a TM-score above 0.7, indicating structural similarity to a known protein fold (Table S5, Figure S11). The majority of designed SEPs thus adopt backbone topologies that are structurally distinct from deposited PDB entries, consistent with their *de novo* origin from RFDiffusion.

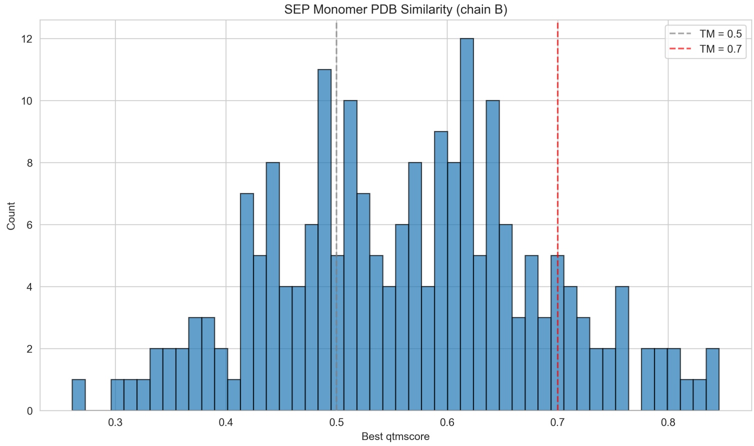

**Figure S11: SEP monomer PDB similarity.** Distribution of query TM-scores for SEP monomer–PDB hits. Dashed lines indicate TM = 0.5 (same fold family) and TM = 0.7 (high structural similarity).

##### C.3.3 VHH–SEP Complex Structural Clustering

**Methods.** The 1,161 VHH–SEP complex structures were clustered using Foldseek `easy-multimercluster` with a TM-score threshold of 0.5 and sensitivity 9.5. This lower threshold (compared to 0.8 for monomers) reflects the greater structural variation expected in binary complexes, where both fold topology and binding geometry contribute to diversity.

**Results.** Clustering yielded **618 unique VHH–SEP complex clusters** (46.8% redundancy; average cluster size 1.9), substantially more diverse than the 278 monomer clusters from the same set of structures (Figure S12). Structural diversity at the complex level indicates that VHH binding mode variation—differences in how SEPs dock against VHH paratopes—contributes substantially to structural diversity beyond SEP fold variation alone.

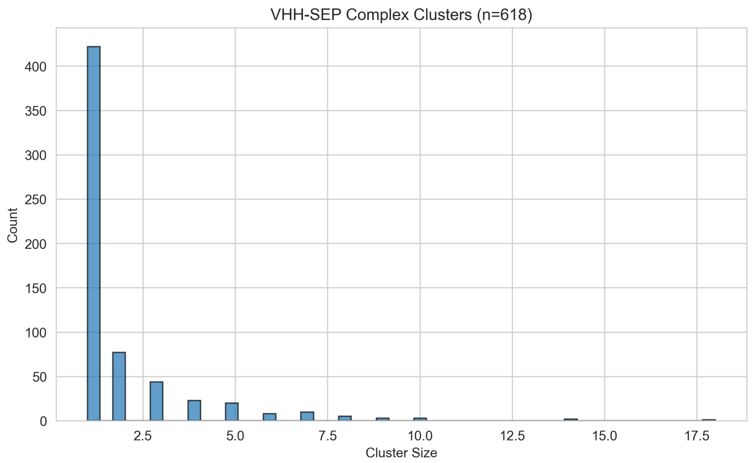

**Figure S12: VHH–SEP complex structural clustering.** Distribution of cluster sizes for 1,161 VHH–SEP complexes clustered at TM-score $\geq 0.5$.

##### C.3.4 VHH–SEP Complex PDB Similarity Search (Multimersearch)

**Methods.** All VHH–SEP complex structures were searched against PDB100 using Foldseek `easy-multimersearch`. To accurately assess whole-complex similarity, we used the complex-level TM-score (`complexqtmscore`) rather than per-chain TM-scores (`qtmscore`), which can yield misleading results for multimers: because per-chain scores are length-weighted averages, a well-aligned VHH chain ($\sim$150 residues) can dominate the score even when the SEP chain ($\sim$50 residues) does not match at all. The `complexqtmscore` evaluates the structural alignment of the entire complex and is normalized by the query complex length, making it robust to target PDB structures with additional chains.

Key search parameters: `cov-mode` 2 (query coverage only, so hits are not penalized for extra chains in PDB assemblies), `min-assigned-chains-ratio` 0.95 (requiring both VHH and SEP chains to be assigned), `coverage` 0.5, `e-value` 1.0, and `max-seqs` 5000.

**Results.** The search returned **15,069 hits** across 2,055 unique query structures. The best PDB match per query had a median `complexqtmscore` of 0.565 and a maximum of 0.728 (Figure S13). While 93.6% of structures had a best match exceeding TM = 0.5 (indicative of the same general fold family, driven largely by the conserved VHH immunoglobulin scaffold), **only 2 of 2,055 structures (0.1%) achieved a `complexqtmscore` above 0.7** (Table S6). Both top hits corresponded to the VHH target 5C2U, which is itself present in the PDB, so the high similarity is expected. No SEPIA structure matched a PDB complex with high whole-complex similarity beyond this trivial case.

**Table S6: Top 5 VHH–SEP complex PDB matches** ranked by `complexqtmscore`. The top hit (5C2U) is the parental VHH structure from the design set.

| VHH Target | PDB Hit | cplxQTM | qCov | E-value |
|---|---|---|---|---|
| 5C2U | 5C2U | 0.728 | 0.984 | $3.5 \times 10^{-24}$ |
| 2VYR | 8J7E | 0.678 | 1.000 | $1.5 \times 10^{-16}$ |
| 2VYR | 8J7E | 0.675 | 1.000 | $1.6 \times 10^{-16}$ |
| 6WAQ | 8J7E | 0.667 | 1.000 | $5.7 \times 10^{-16}$ |
| 6WAQ | 5BOP | 0.665 | 0.976 | $3.1 \times 10^{-18}$ |

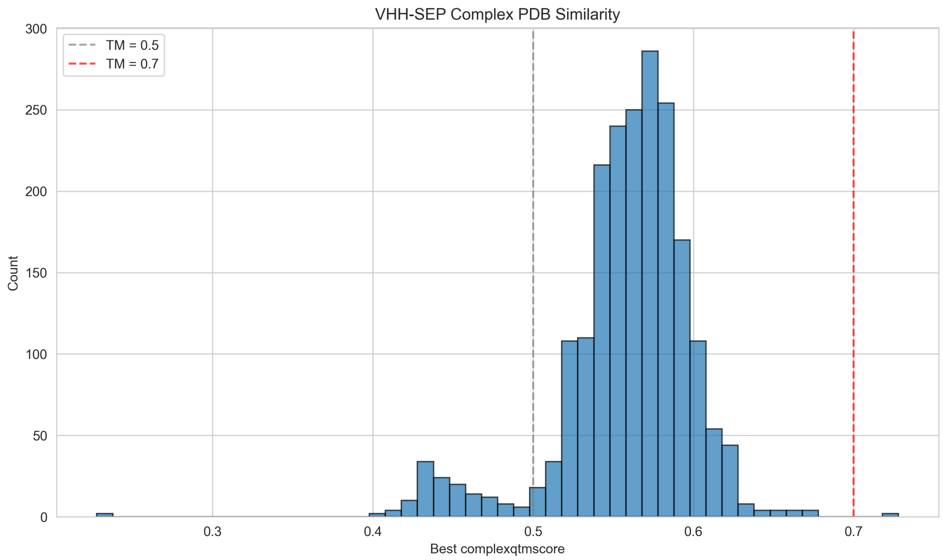

**Figure S13: VHH–SEP complex PDB similarity.** Distribution of complex-level query TM-scores for PDB hits from Foldseek multimersearch. Dashed lines indicate TM = 0.5 and TM = 0.7.

**VHH–SEP complex comparison to PDB.** To assess whether the designed VHH–SEP complexes structurally resemble known VHH–antigen complexes in the PDB, we used Foldseek multimersearch to find the closest PDB match for each of 78 unique VHH targets with at least one validated SEP hit across Rounds 1 and 2. Table S7 shows the 20 VHH–SEP complexes with the highest structural similarity to any PDB entry. Even among the top matches, only one VHH–SEP complex (5C2U) exceeds TM = 0.7, and this is a trivial self-match where the closest PDB hit is the native VHH–antigen crystal structure from which the VHH was derived. The median closest-match score across all 78 targets is 0.583 (well below the TM = 0.7 threshold for high structural similarity), demonstrating that VHH–SEP complexes adopt binding geometries distinct from the VHH–antigen complexes deposited in the PDB.

**Table S7: Structural comparison of VHH–SEP complexes to the PDB.** Top 20 VHH–SEP complexes ranked by structural similarity (`complexqtmscore`) to their closest match in PDB100 via Foldseek multimersearch. Despite sharing the same VHH scaffold, designed VHH–SEP complexes show low structural similarity to known VHH–antigen complexes, with only one exceeding TMscore = 0.7. Self-match indicates the closest PDB hit is the native VHH–antigen crystal structure from which the VHH was derived.

| VHH Target | Closest PDB Hit | cplxQTM | qCov | Self-match |
|---|---|---|---|---|
| 5C2U | 5C2U | 0.728 | 0.903 | Yes |
| 2VYR | 8J7E | 0.678 | 1.000 | |
| 6WAQ | 8J7E | 0.667 | 1.000 | |
| 6U54 | 7C6A | 0.656 | 1.000 | |
| 5MP2 | 5F1K | 0.651 | 0.992 | |
| 6U51 | 8J7E | 0.643 | 0.873 | |
| 5VXM | 5VXM | 0.634 | 0.662 | Yes |
| 8DQU | 4N9O | 0.626 | 1.000 | |
| 8DAM | 4WEU | 0.622 | 0.507 | |
| 5VXL | 5VXL | 0.620 | 1.000 | Yes |
| 4NC1 | 5VXM | 0.619 | 0.557 | |
| 5VXK | 5VXK | 0.614 | 0.545 | Yes |
| 5BOP | 4U0R | 0.612 | 0.566 | |
| 5M2J | 6FUZ | 0.611 | 0.974 | |
| 5IMM | 5VXK | 0.605 | 0.589 | |
| 6XZU | 4YGA | 0.604 | 1.000 | |
| 5M2I | 6YU8 | 0.603 | 0.521 | |
| 5FV2 | 7SO7 | 0.602 | 0.500 | |
| 6Z1V | 5VXM | 0.601 | 0.506 | |
| 6LZ2 | 7C6A | 0.601 | 0.966 | |

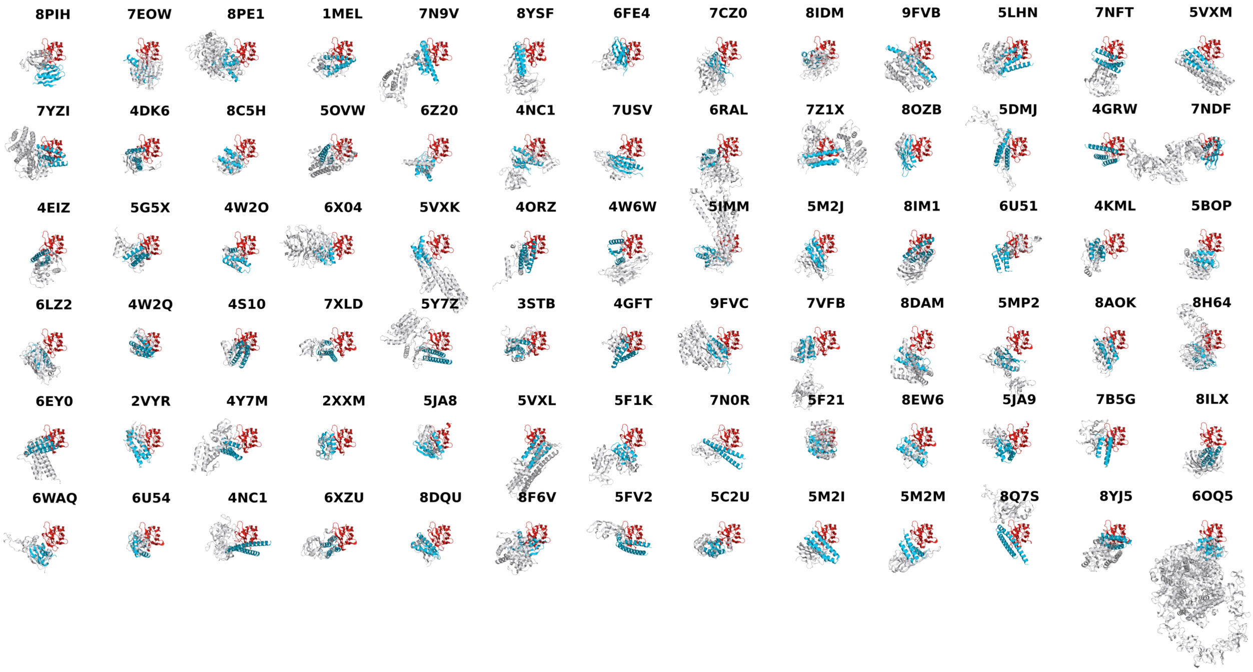

**Figure S14: Structural overlay of VHH–SEP complexes with native VHH–antigen structures.** For each of 78 unique VHHs targeted by SEPIA (Rounds 1 and 2 combined, deduplicated by VHH sequence), the highest-affinity SEP hit is shown overlaid with the native VHH–antigen complex from the PDB. VHH chains (red) are aligned across all structures; SEPs are shown in blue and native antigens in grey. PDB identifiers for the native VHH–antigen structures are shown above each panel. Structures are ordered by SEP binding affinity (strongest top-left to weakest bottom-right).

**Structural overlay of VHH–SEP complexes with native VHH–antigen structures.** For each of the 78 unique wild-type VHH sequences with at least one validated SEP hit, we superimposed the highest-affinity predicted VHH–SEP complex structure with the native VHH–antigen crystal structure from the PDB, aligned on the VHH chain (Figure S14). Structures are arranged by binding affinity (strongest first, left to right, top to bottom). Consistent with the Foldseek structural similarity analyses above (Table S8, Figure S13, Table S7), SEPs (blue) are structurally distinct from native antigens (grey) and frequently engage different paratope regions on the VHH (red) than those contacted by the native antigen.

**Summary: multimer vs. monomer comparison.** Table S8 summarizes the key differences between SEP monomer and VHH–SEP complex structural analyses.

**Table S8: Comparison of SEP monomer vs. VHH–SEP complex structural analysis.**

| Metric | VHH–SEP Complex | SEP Monomer |
|---|---|---|
| Input structures | 1,161 | 1,161 |
| Unique clusters | 618 | 278 |
| Clustering TM-score threshold | 0.5 | 0.8 |
| Clustering redundancy | 46.8% | 76.1% |
| Total PDB hits | 15,069 | 4,367 |
| Structures with PDB hits | 2,055 | 205 |
| Median best PDB TM-score | 0.565 | 0.568 |
| Structures with TM-score > 0.7 | 2 (0.1%) | 26 (12.7%) |

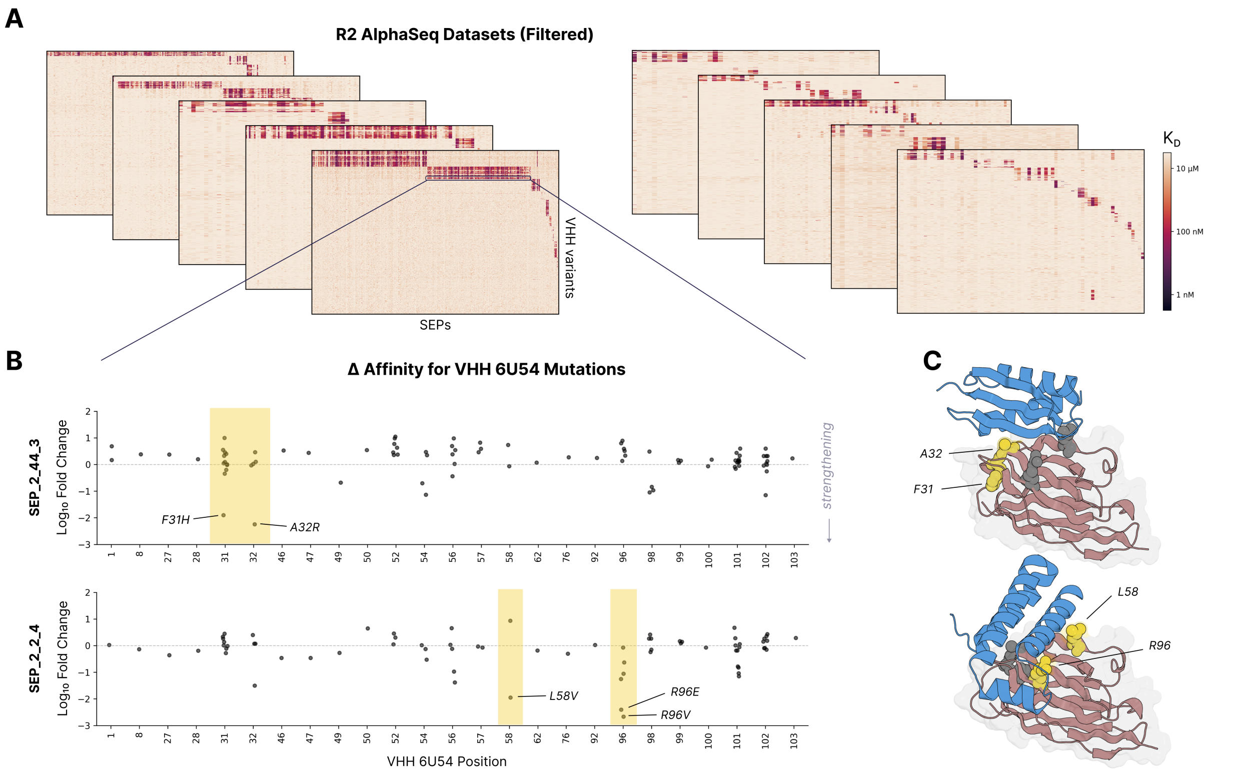

**Figure S15: Round 2 AlphaSeq datasets and VHH mutational analysis. (A)** Filtered AlphaSeq affinity heatmaps for Round 2 experiments. Each heatmap represents a single AlphaSeq experiment, showing VHH variants (rows) against SEPs (columns) for a different VHH target. Color scale indicates AlphaSeq binding affinity strength (darker = stronger binding). **(B)** Change in AlphaSeq affinity for VHH point mutations in VHH 6U54, measured against two representative SEPs (SEP_2_44_3, top; SEP_2_2_4, bottom). Yellow indicates positions that improve binding affinity. Binding-enhancing mutations are labeled. Negative fold change = mutation improves binding affinity. **(C)** Structural mapping of key mutation sites (yellow spheres) onto VHH 6U54–SEP pseudo-structures, showing that affinity-sensitive residues (F31, A32 for SEP_2_44_3, top; L58, R96 for SEP_2_2_4, bottom) localize to the predicted binding interface.

#### C.4 VHH–SEP Residue Coupling Analysis

This section presents residue-level analyses of VHH–SEP binding interfaces from Round 2 VHH mutational landscapes and Round 3 targeted mutational scans.

##### C.4.1 Round 2 VHH Mutational Landscapes

Round 2 included 11,425 VHH point mutants targeting 180 parental VHHs, enabling systematic analysis of how single-residue changes at the paratope affect SEP binding specificity. Figure S15A shows filtered AlphaSeq affinity heatmaps for Round 2 experiments, with each heatmap representing VHH variants (rows) screened against SEPs (columns) for a single VHH target.

Figure S15B illustrates the effect of VHH point mutations on binding affinity for VHH 6U54 against two representative SEPs. Mutations at specific positions substantially enhance or ablate binding, and the identity of these positions differs between SEPs, consistent with each SEP engaging a distinct subset of paratope residues. Structural mapping of the most affinity-sensitive positions onto the designed VHH–SEP pseudo-structures (Figure S15C) confirms that these residues localize to the predicted binding interface. Together, these data reveal a rich landscape of tolerated and deleterious mutations at VHH–SEP interfaces, substantially increasing the scale and granularity of joint sequence–structure–affinity data within SEPIA.

##### C.4.2 Round 3 Alanine Scans

To validate whether the designed synthetic epitopes form structurally accurate contacts with their cognate VHHs, we analyzed single-residue perturbation data across 96 VHH–SEP pairs spanning 5 parent VHHs. This analysis combines double-sided alanine scanning data from 4 parent VHHs (4NC1, 5M2I, 6XZU, and 6U54; Section B.4.3) with site saturation mutagenesis (SSM) data from 7XLD (Section C.4.3), each parent VHH paired with multiple SEP variants. We measured binding affinities using AlphaSeq and computed the change in affinity ($\Delta K_d$) resulting from each single-residue perturbation relative to the wild-type SEP.

We evaluated whether $\Delta K_d$ values could discriminate interface residues from non-interface residues by computing the AUROC with respect to a binary contact label, defined as whether the perturbed residue's C$\alpha$ atom lies within 8 Å of any VHH residue in the predicted complex structure. Missing $\Delta K_d$ values, which typically indicate mutations that ablated binding below the assay's detection threshold, were imputed to the 95th percentile of observed values within each VHH–SEP pair.

We find that 88 of 96 VHH–SEP pairs (91%) exhibit better-than-random AUROC for contact prediction. At the pooled level, $\Delta K_d$ achieves an AUROC of 0.63, with contact residues showing significantly elevated $\Delta K_d$ compared to non-contact residues (Mann-Whitney $U$ test, $p < 10^{-30}$). Notably, contact positions exhibited a 38% higher rate of missing (unmeasurable) binding values compared to non-contact positions (34% vs. 25%, $\chi^2$ test $p < 10^{-100}$), further supporting that interface residues are functionally critical for binding. This function-to-structure concordance provides orthogonal validation that the designed SEPs engage their target VHHs through the intended structural interfaces.

##### C.4.3 Round 3 SEP Site Saturating Mutagenesis

To systematically characterize the sequence determinants of SEP binding, we performed single-site saturation mutagenesis (SSM) on validated SEP designs targeting the 7XLD_B parent VHH.

**Library design.** For a parental SEP, we generated variants with single amino acid substitutions at every position to all 18 alternative amino acids (excluding cysteine). Additionally, we included random pair mutations to sample epistatic interactions. The complete library comprised 4,779 SEP variants and the WT. Our analysis is limited to the single-variants to demonstrate sensitivity of the paratope/epitope interaction.

**Results.** Of 4,779 SEP variants screened against wild-type 7XLD_B, 3,496 variants (73.2%) met our hit criteria. Analysis of mutational tolerance revealed that positions predicted *in silico* to contact the VHH were significantly less tolerant to mutation than non-contact positions, consistent with their functional importance for binding. Conversely, solvent-exposed positions distant from the interface were generally more tolerant to substitution. Substitutions to proline (P) are deleterious at most residues, likely indicative of breaking proper protein folding for the SEP.

### D Evaluation Datasets for ML Models

#### D.1 SAbDab-nano Decoy Dataset

We constructed a decoy dataset from SAbDab-nano to evaluate model discrimination between true binders and structurally plausible non-binders.

**Source data selection.** We extracted all VHH–antigen complexes from SAbDab-nano and applied the following filtering criteria:

- Retained only complexes where antigen type is "protein" (excluding peptides, haptens, and multi-chain antigen complexes)
- Cleaned VHH sequences using ANARCI to remove non-standard residues and ensure consistent numbering
- Applied an antigen length cap of $\sim$500 residues

**On-target complex generation.** On-target interactions are defined as VHH–antigen pairs with solved co-crystal structures in SAbDab-nano, which are by definition experimentally validated binders. For each pair, we generated three Boltz-2 complex predictions using the known PDB co-structure as a structural template. No MSAs were used during prediction. We selected the prediction with highest ipSAE score as the representative on-target structure.

**Off-target (decoy) complex generation.** Decoy complexes were generated for mismatched VHH–antigen pairs previously validated as non-interacting in AlphaSeq. To ensure consistent templating between on-target and off-target predictions, we used a two-stage approach: first, three Boltz-2 predictions were generated per pair using AlphaFold-predicted antigen monomer templates; the best Stage 1 prediction (by TM-score to template) was then used as a complex template for a second round of three Boltz-2 predictions. The Stage 2 prediction with the highest ipSAE score was selected as the representative decoy structure.

**Dataset statistics.** After prediction and filtering, the final benchmark dataset consists of:

- 646 on-target complexes (labeled "bind")
- 14,521 off-target decoy complexes (labeled "non-bind")
- ∼38.8 decoys per antigen on average
- Decoy ratio: 22.5:1 (off-target:on-target)

Dataset composition and structural quality metrics are summarized in Table S9 and Figures S16–S17 and Figure S18. MICAN TM-scores are reported separately for on-target predictions (Figure S18), as this metric requires experimental reference structures unavailable for off-target decoys; the high median TM-score (0.942) confirms that predicted on-target complex structures closely recapitulate the crystallographic binding modes.

**Table S9:** SAbDab-nano decoy dataset composition and quality metrics

| Metric | On-Target | Off-Target | Total |
|---|---|---|---|
| Total PPIs | 646 | 14,521 | 15,167 |
| Unique VHH | 595 | 960 | 960 |
| Unique Antigens | 251 | 374 | 438 |
| Decoy ratio | – | 22.5:1 | – |
| *Quality Metrics (median values)* | | | |
| ipSAE | 0.81 | 0.28 | – |
| pTM complex | 0.92 | 0.85 | – |
| iPTM complex | 0.95 | 0.84 | – |
| pLDDT complex | 0.89 | 0.84 | – |
| MICAN TM-score[a] | 0.942 | – | – |

[a]MICAN TM-score only computed for on-target predictions against experimental structures; not applicable to off-target decoys which lack experimental complex structures.

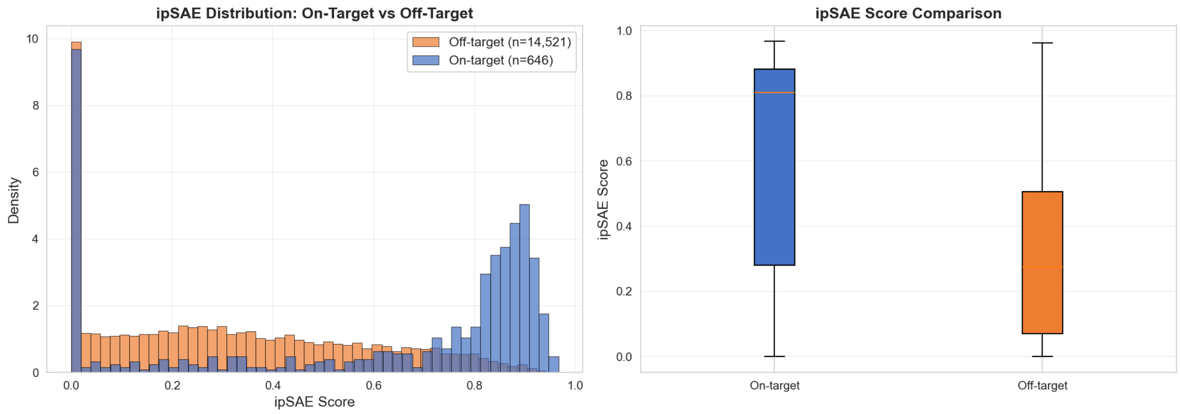

**Figure S16: ipSAE score distribution for SAbDab-nano decoy dataset. Left**: Histogram showing on-target (blue, n=646) and off-target (coral, n=14,521) ipSAE score distributions. On-target predictions have significantly higher ipSAE scores (median 0.81 vs 0.28, 2.93× fold change). **Right**: Box plot comparison showing clear separation between on-target and off-target predictions.

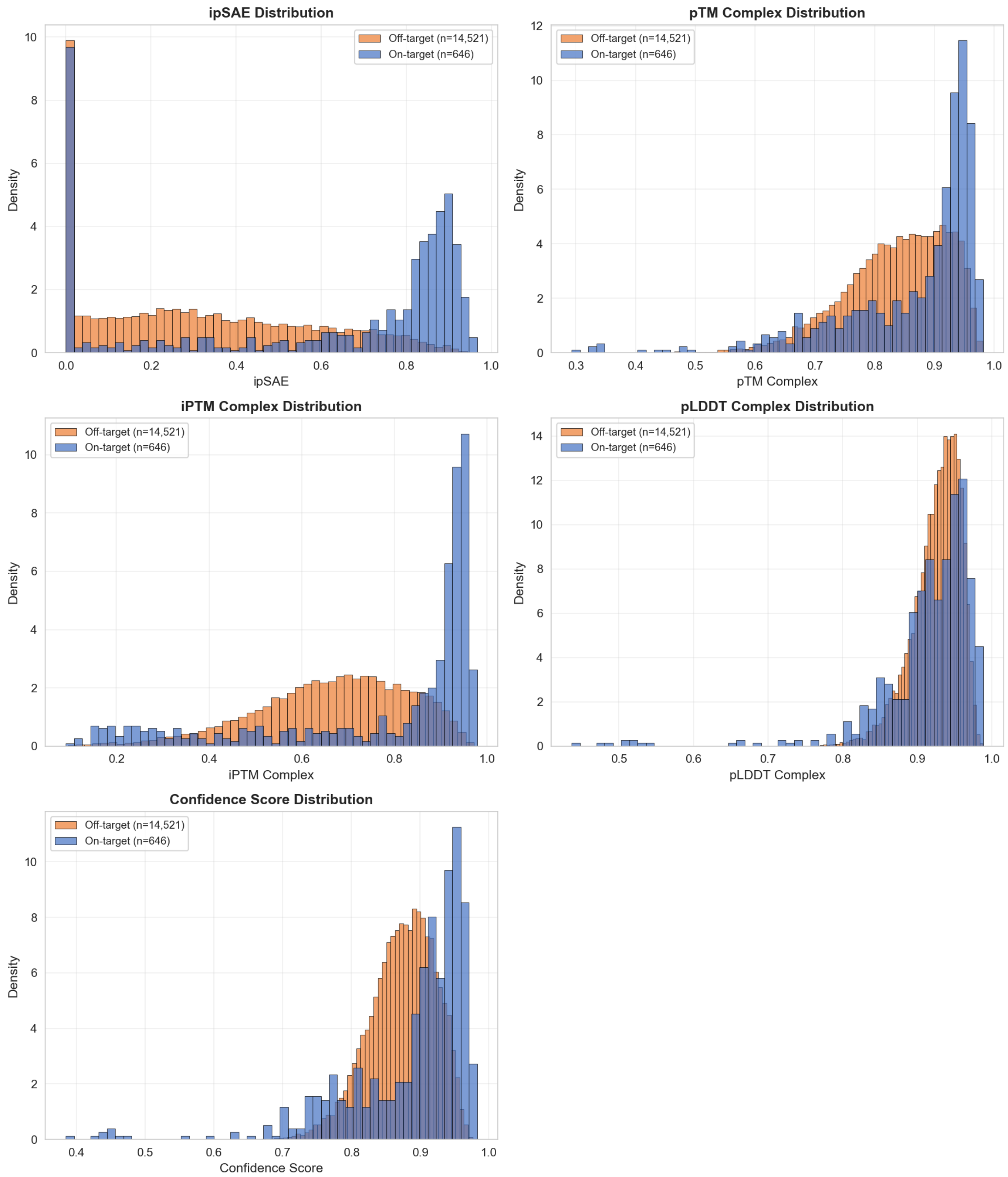

**Figure S17: Multi-metric comparison for SAbDab-nano decoy dataset.** Distribution of five quality metrics comparing on-target (blue, n=646) and off-target (coral, n=14,521) predictions. All metrics show consistent trends with on-target predictions having higher scores, with ipSAE showing the strongest separation (2.93× fold change in medians). MICAN TM-scores are only computed for on-target predictions (see separate figure) as off-target predictions lack experimental reference structures.

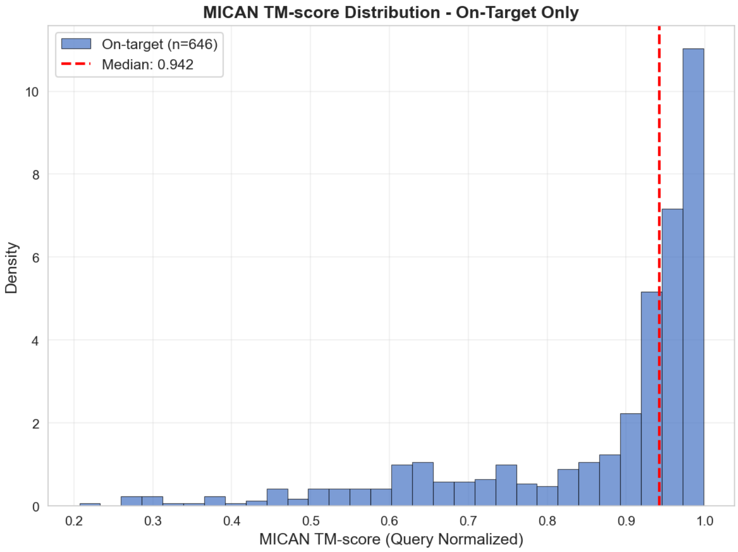

**Figure S18: MICAN TM-score for on-target predictions.** Density distribution of MICAN TM-scores (query-normalized) for on-target predictions assessed against experimental crystal structures from SAbDab-nano. Median TM-score: 0.942 (dashed red line). MICAN TM-scores are not computed for off-target predictions as these non-binding pairs lack experimental complex structures.

#### D.2 De Novo VHH Dataset

To evaluate ML model generalization beyond SEPIA VHH–SEP structures, we assembled a *de novo* VHH–antigen dataset comprising VHH designs generated using open-source *de novo* antibody design methods and tested against cognate targets in AlphaSeq. This dataset serves as an out-of-distribution evaluation set: VHH sequences were generated *de novo* (not from PDB structures), target antigens have no known VHH binders in training data, experimental validation provides ground-truth labels, and the low hit rate creates challenging class imbalance.

**Antigen selection.** We selected 22 antigen targets from recent *de novo* antibody design studies [13–15], none of which have an antibody-bound co-crystal structure in the PDB (Table S10), requiring *de novo* design to operate without direct structural precedent for antibody binding for all targets.

**Experimental validation.** 4,930 candidate VHHs (generated using several open-source *de novo* VHH design methods) were measured via AlphaSeq. Applying affinity-based hit criteria (Section B.3), we identified 37 VHH–antigen hits (0.8% VHH hit rate), spanning 8 of 22 antigens (36.4%). A per-antigen breakdown of candidates and hits is provided in Table S10.

##### D.2.1 *De Novo* VHH Structural Analysis

We characterized the structural relationship between *de novo* VHH–antigen complexes, the PDB, and SEPIA VHH–SEP structures using Foldseek [4].

**Similarity to the PDB.** We compared all 37 validated *de novo* VHH–antigen complexes against the PDB using Foldseek `multimersearch`. We evaluated complex-level TM-scores (`complexqtmscore`), which capture whole-complex similarity rather than per-chain averages that can be dominated by the conserved VHH scaffold. The median best-match TM-score was 0.455 (below the same-fold threshold of 0.5), and no complexes exceeded 0.7 (Figure S19).

**Structural comparison to SEPIA.** We pooled 1,161 SEPIA VHH–SEP hit complexes with the 37 *de novo* hit complexes and clustered them with Foldseek `multimercluster` (TM-score threshold 0.5). Of 628 resulting clusters, 619 contained only SEPIA structures and 9 contained only *de novo* structures; no cluster contained members from both datasets (Table S11). At the monomer level, clustering SEP chains against antigen chains (TM-score threshold 0.8) yielded 287 clusters with zero mixed membership. These results indicate that *de novo* VHH–antigen complexes occupy distinct structural space from both the PDB and SEPIA VHH–SEP structures.

**Table S10:** Per-antigen breakdown of the *de novo* VHH dataset. Hit criteria are defined in Section B.3 and Table S1.

| Antigen | UniProt ID | Candidates | Hits | Hit Rate (%) |
|---|---|---|---|---|
| RFK | Q969G6 | 194 | 12 | 6.2 |
| BHRF1_viral | P03182 | 200 | 8 | 4.0 |
| ORM2 | P19652 | 271 | 7 | 2.6 |
| IFNAR2 | P48551 | 234 | 2 | 0.9 |
| IL3 | P08700 | 384 | 3 | 0.8 |
| CEACAM6 | P40199 | 299 | 2 | 0.7 |
| IL10Ra | Q13651 | 398 | 2 | 0.5 |
| CRLF2 | Q9HC73 | 350 | 1 | 0.3 |
| CD2 | P06729 | 33 | 0 | 0.0 |
| CEACAM1 | P13688 | 395 | 0 | 0.0 |
| CR2 | P20023 | 94 | 0 | 0.0 |
| GCSFR_CSF3R | Q99062 | 138 | 0 | 0.0 |
| GCSF_CSF3 | P09919 | 120 | 0 | 0.0 |
| IL13Ra2 | Q14627 | 2 | 0 | 0.0 |
| IL15Ra | Q13261 | 155 | 0 | 0.0 |
| IL20 | Q9NYY1 | 279 | 0 | 0.0 |
| IL6Rb | P40189 | 170 | 0 | 0.0 |
| IL9R | Q01113 | 176 | 0 | 0.0 |
| MZB1 | Q8WU39 | 61 | 0 | 0.0 |
| OncostatinM | P13725 | 183 | 0 | 0.0 |
| PMVK | Q15126 | 398 | 0 | 0.0 |
| somatotropin | P01241 | 396 | 0 | 0.0 |
| **Total** | | **4,930** | **37** | **0.8** |

8 of 22 antigens yielded at least one hit. None of the 22 antigens have an antibody-bound co-crystal structure in the PDB.

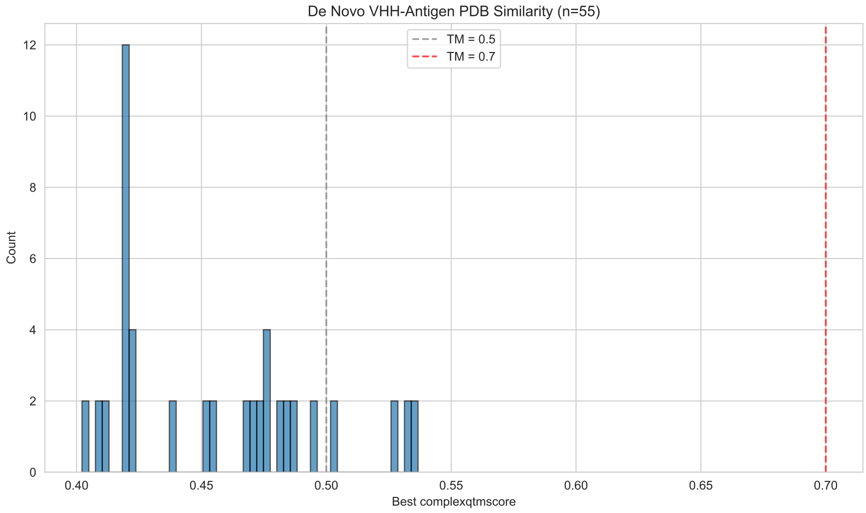

**Figure S19: Structural similarity of *de novo* VHH–antigen complexes to the PDB.** Distribution of best complex-level TM-scores (`complexqtmscore`) from Foldseek `multimersearch` against the PDB for 37 validated *de novo* VHH–antigen complexes. Median best-match TM-score: 0.455 (below the same-fold threshold of 0.5). No complexes exceed TM = 0.7, confirming that all *de novo* complexes occupy novel structural space relative to the PDB.

**Table S11:** Structural clustering of SEPIA and *de novo* VHH datasets. Clustering performed using Foldseek at TM-score threshold of 0.5 for complexes and 0.8 for monomers. Mixed clusters contain structures from both datasets.

| **Analysis** | **Total Clusters** | **SEPIA-only** | ***De novo*-only** | **Mixed** |
|---|---|---|---|---|
| VHH-epitope complexes | 628 | 619 (98.6%) | 9 (1.4%) | 0 (0.0%) |
| Epitope monomers | 287 | 281 (97.9%) | 6 (2.1%) | 0 (0.0%) |

**Table S12:** Examples of detected UniRef50 outlier sequences.

| UniProt ID | Amino Acid Sequence |
|---|---|
| Q156A1 | MQQQQQQQQQQQQQQQQQQQQQQQQQQQQQQQQQQQQQQQQQQQ QQQQQQQQQQQQQQQQQQQQQQQQQQQQQQQQQQQQQQQ |
| A0A091WA51 | GRGREGKGKERKGKERKGKERKGKERKGKERKGKERKGKERKGKE RKGKERKGKERK |
| UPI00211E4217 | DSDSDSDSDSDSDSDSDSDSDSDSDSDSDSDSDSDSDSDSDSDSDSDS DSDSDSDSDSDSDSDSDSDSDSDSDSDSDSDSDSDSDSDSDSDSDSDS DSDSDSDSDSDSDSDSDSDSDSDGD |
| UPI001122B86D | MLVLVLVLVLVLVLVLVLVLVLVLVLVLVLVLVLVLVLVLVLVLVL CRRSDAAEPRARDVEVAAPGKSSCGAR |

### E ML Models

#### E.1 Sequence Outlier Detector

We trained an outlier detector to filter unlikely amino acid sequences from our SEPIA *de novo* design pipelines. We used UniRef50 [16, 17] (downloaded March 2025) as training data, representing a distribution of natural and likely functional protein sequences. We filtered to UniRef50 sequences in the length range of 50–500 amino acids, yielding $\sim$500,000 unique sequences.

Protein sequences were featurized using feature hashing [18] of amino acid trigrams into 43 buckets, producing a fixed-length count vector per sequence.

We trained an isolation forest [19] as implemented in scikit-learn [20] with default hyperparameters (100 estimators) on the full encoded UniRef50 set. The outlier threshold was set to $t < -0.05$. Table S12 lists examples of UniRef50 outliers detected by the model.

#### E.2 Neighborhood Transformer (NTX)

We designed and trained the Neighborhood Transformer (NTX), a structure-based model that controls for data leakage of previously seen protein–protein complexes. The NTX training distribution contains no protein–protein complexes or antibody structures to enable fair evaluation on the decoy dataset.

NTX is a protein geometric language model that combines concepts from protein language models, inverse folding, and graph neural networks. As described below, NTX is trained with a masked language model loss analogous to BERT-style transformers, comparable to the Masked Inverse Folding (MIF) [21] and ESM-IF [22] models. However, the attention layers are restricted to only residues within a local neighborhood (12 Å radius) of each central masked residue. Attention is further augmented by incorporating geometry of the local neighborhood as edge features.

##### E.2.1 Feature Inputs

NTX operates at backbone-only resolution. For each $i$-th residue, the Cartesian coordinates of backbone atoms $C_i$, $N_i$, $O_i$, $C\alpha_i$, and a virtual $C\beta_i$ (inferred as in RFDiffusion [3]) are extracted. A neighborhood is defined as the $k \leq 32$ nearest residues within a 12 Å $C\alpha$–$C\alpha$ radius.

SE(3)-invariant node and edge features are computed for each neighborhood: (1) Gaussian-kernel for pairwise distances $r_{ij}^{ab}$ between each backbone atom $a$ (belonging to the central residue $i$) and backbone each atom $b$ (belonging to neighbor residue $j$), $g_{ij}^{ab} = \exp[-0.5\,(r_{ij}^{ab}/\sigma)^2]$ with $\sigma$=4 Å, plus inter-neighbor distances $g_{jk}^{bc}$; (2) dot products of backbone normal vectors $\phi_{ij}^{ab} = \vec{v}_i^{\,a} \cdot \vec{v}_j^{\,b}$, with $\vec{v}_i^{\,a} = (\vec{r}_i^{\,a} - \vec{r}_i^{\,C_\alpha})/||\vec{r}_i^{\,a} - \vec{r}_i^{\,C_\alpha}||$; (3) a per-residue SASA feature computed from

110-point Lebedev quadrature grids (radius 4 Å) centered at C$\alpha_i$, rotated into the local residue frame and encoded as binary solvent-exposure indicators; and (4) one-hot amino acid identities of all $j$ neighbors (the $i$-th residue identity is masked).

###### E.2.2 Architecture

Node and edge features are projected to a $d$=500 latent space via per-type linear layers. Central-residue features are broadcast and summed with each neighbor's features, forming a (32x500) equivariant representation (no positional encoding; zero-filled and attention-masked for neighborhoods with fewer than 32 residues).

This representation passes through 6 edge-aware transformer encoder blocks (SwiGLU activations, pre-norm layer normalization, 4 attention heads). Edge features are encoded per layer by a shallow MLP into a 4-dimensional bias $B_{jk}$ added respectively to the 4 attention head logits: $\mathrm{Attn}(Q, K, V) = \mathrm{softmax}(L + B)\,V$ with $L_{jk} = Q_j K_k/\sqrt{d}$. After the transformer blocks, neighbor representations are aggregated via a distance-weighted sum (weights $g_{ij}^{C\alpha,C\alpha}$), yielding a 500-dimensional latent vector per residue. A final two-layer MLP (250-dimensional hidden layer, SwiGLU, then log-softmax over 20 amino acids) produces the masked-residue prediction. NTX contains approximately 30 million trainable parameters.

###### E.2.3 Pre-training

NTX was pre-trained on monomer structures from the AlphaFold Protein Structure Database (AFDB) [23], which contains no protein complexes. We selected Swiss-Prot–annotated entries plus model organism and global health proteomes, then removed any structures present in the ProteinMPNN [5] training/test sets (by sequence or ID match) as well as all antibody structures from SAbDab [9], yielding 782,506 unique AFDB structures (752,506 training, 30,000 validation).

Training minimized a masked language model cross-entropy loss over 20 amino acid identities using the Adam optimizer (learning rate $10^{-4}$, halved after 10 epochs without validation improvement). Each epoch comprised 4,000 minibatches of sampling from 500 structures each, with one residue per structure sampled via approximate stratification over amino acid identities. Gaussian noise $\mathcal{N}(0, 0.025\text{ Å})$ was applied to atomic coordinates as data augmentation. Training converged at $\sim$73% validation accuracy after $\sim$24 hours on an Nvidia L40S GPU.

###### E.2.4 Embeddings

NTX produces a 500-dimensional latent vector per residue. For a protein of length $L$, running NTX over all residues yields an ($L$, 500) structure-aware embedding $Z$.

To capture binding-specific signal, we compute confidence-weighted embedding differences between the complex and its constituent monomers (embedded in their holo conformations):

$$Z_i = p^a_{i,\mathrm{complex}} Z_{i,\mathrm{complex}} - p^a_{i,\mathrm{monomer}} Z_{i,\mathrm{monomer}}$$

where $p^a_i$ is the NTX confidence for the known amino acid $a$ at residue $i$. $Z_i$ is zero for residues whose neighborhood is unchanged between monomer and complex, naturally restricting the representation to interfacial residues. For downstream ML models, we aggregate over residues: $Z = \sum_i Z_i$.

##### E.3 ABACUS

Here we discuss the details of our ABACUS classifier.

**Embedding source.** For each predicted complex structure, we extracted embeddings from the final layer of the Boltz-2 trunk representation including the single representations $\mathbf{s} \in \mathbb{R}^{N\times d_s}$ and pair representations $\mathbf{z} \in \mathbb{R}^{N\times N\times d_z}$, where $N$ is the total number of residues in the complex. In preliminary experiments, trunk embeddings yielded higher performance than confidence module embeddings, despite interface confidence scores (such as iPTM or ipSAE) being a standard method for filtering binders from non-binders. This may be due to richer information likely present in the trunk embeddings, as the final confidence embeddings need to solely contain the information necessary for predicting the final PAE scores, whereas the trunk embeddings must encode the information required for complex generation, and ultimately the final confidence as well, while still having gone through significant amount of non-linear processing through the previous trunk layers.

**Interface definition.** The antibody-antigen interface is defined using a 10 Å C$\alpha$–C$\alpha$ distance threshold:

- **Paratope**: The set of antibody (VHH) residues with at least one C$\alpha$ atom within 10 Å of any antigen C$\alpha$ atom
- **Epitope**: The set of antigen residues with at least one C$\alpha$ atom within 10 Å of any antibody C$\alpha$ atom

Interface residue positions are pre-computed from the predicted 3D structures and stored as position indices.

**Single-residue embeddings (s).** For each interface residue $i$, we extract its $d_s$-dimensional embedding from the full residue representation. The paratope and epitope embeddings are concatenated:

$$\mathbf{s} = [\mathbf{s}_{\text{para}}; \mathbf{s}_{\text{epi}}] \in \mathbb{R}^{(n_{\text{para}}+n_{\text{epi}})\times d_s} \tag{1}$$

**Pairwise embeddings (z).** We extract cross-interface pairwise representations $\mathbf{z}_{\text{para}\to\text{epi}} \in \mathbb{R}^{n_{\text{para}}\times n_{\text{epi}}\times d_z}$ encoding the interaction context between each paratope-epitope residue pair. To capture bidirectional interactions, we additionally extract the reverse direction $\mathbf{z}_{\text{epi}\to\text{para}} \in \mathbb{R}^{n_{\text{epi}}\times n_{\text{para}}\times d_z}$, enabling symmetric processing of the interface.

**Predicted Aligned Error (PAE) features.** We augment embeddings with PAE-derived confidence features that provide information of the Boltz-2 model's confidence at individual residues.

*For single-residue embeddings:* For each interface residue $i$, we compute aggregated PAE statistics capturing both local prediction confidence and cross-interface uncertainty:

- **Intra-region PAE**: Mean, minimum, and maximum PAE values between residue $i$ and other residues in the same region (paratope or epitope), excluding self-interactions. These statistics reflect local structural confidence within each binding partner.
- **Cross-interface PAE**: Mean, minimum, and maximum PAE values between residue $i$ and all residues in the opposite region. These statistics capture the model's confidence in the relative positioning across the interface, which is particularly informative for binding prediction.

This yields 6 additional features per residue (3 aggregations $\times$ 2 contexts):

$$\mathbf{s}_i^{\text{PAE}} = [\mu_{\text{intra}}, \min_{\text{intra}}, \max_{\text{intra}}, \mu_{\text{cross}}, \min_{\text{cross}}, \max_{\text{cross}}] \tag{2}$$

*For pairwise embeddings:* The PAE matrix is asymmetric: PAE$[i, j]$ represents the predicted error in residue $i$'s position when the prediction is aligned to residue $j$. For each residue pair $(i, j)$ in the cross-interface submatrix, we concatenate the corresponding PAE value as an additional feature:

$$\mathbf{z}'_{ij} = [\mathbf{z}_{ij}; \text{PAE}[i, j]] \in \mathbb{R}^{d_z+1} \tag{3}$$

**Global confidence features.** We incorporate 8 structure prediction confidence metrics as global features: confidence score, pTM, ipTM, protein ipTM, complex pLDDT, complex ipLDDT, complex PDE, and complex iPDE. These metrics are extracted from the Boltz-2 confidence output for each predicted structure and provide complementary global assessments of prediction quality.

The global confidence vector $\mathbf{c} \in \mathbb{R}^{d_c}$ is concatenated to the final pooled representation before the classification head, providing a direct signal about overall prediction quality to the classifier.

##### E.3.1 Per-Residue MLP

The PAE-augmented single-residue embeddings $\mathbf{s}' \in \mathbb{R}^{(d_s+6)}$ are processed by a two-layer MLP to project them into a task-specific representation space:

$$\mathbf{h} = \text{MLP}_s(\mathbf{s}') = \text{Dropout}\left(\text{ReLU}\left(\mathbf{W}_2 \cdot \text{Dropout}\left(\text{ReLU}\left(\mathbf{W}_1\mathbf{s}' + \mathbf{b}_1\right)\right) + \mathbf{b}_2\right)\right) \tag{4}$$

where $\mathbf{W}_1 \in \mathbb{R}^{64\times(d_s+6)}$, $\mathbf{W}_2 \in \mathbb{R}^{32\times 64}$, and dropout probability $p = 0.2$.

##### E.3.2 Pairwise Embedding Processing

The PAE-augmented pairwise embeddings $\mathbf{z}' \in \mathbb{R}^{(d_z+1)}$ undergo analogous transformation via a dedicated MLP:

$$\mathbf{z}'' = \text{MLP}_z(\mathbf{z}') = \text{Dropout}\left(\text{ReLU}\left(\mathbf{W}_4 \cdot \text{Dropout}\left(\text{ReLU}\left(\mathbf{W}_3\mathbf{z}' + \mathbf{b}_3\right)\right) + \mathbf{b}_4\right)\right) \tag{5}$$

with $\mathbf{W}_3 \in \mathbb{R}^{64\times(d_z+1)}$ and $\mathbf{W}_4 \in \mathbb{R}^{32\times 64}$.

###### E.3.3 Pairwise Aggregation via Axial Attention

To aggregate pairwise information, we employ axial attention that factorizes attention over the 2D residue-pair matrix. In preliminary experiments, this design achieved better performance than a full pairformer architecture with triangular attention updates while being substantially faster to train.

**Row attention.** For each paratope position $i$, we apply multi-head self-attention across all epitope positions, allowing the model to learn which epitope residues are most relevant for each paratope residue:

$$\mathbf{z}'_{\text{row}}[i,:] = \text{TransformerEncoder}(\mathbf{z}'[i,:]) \quad \text{for } i = 1, \ldots, n_{\text{para}} \tag{6}$$

**Column attention.** For each epitope position $j$, we apply multi-head self-attention across all paratope positions:

$$\mathbf{z}''[:,j] = \text{TransformerEncoder}(\mathbf{z}'_{\text{row}}[:,j]) \quad \text{for } j = 1, \ldots, n_{\text{epi}} \tag{7}$$

Each transformer encoder consists of a single layer with 4 attention heads, hidden dimension 32, feedforward dimension 64, and dropout 0.1.

**Symmetric processing.** The pairwise representation $\mathbf{z}$ is inherently directional: $\mathbf{z}[i,j]$ encodes information about the relationship from residue $i$ to residue $j$, which may differ from $\mathbf{z}[j,i]$. To capture both directions of the antibody-antigen interaction, we process both $\mathbf{z}_{\text{para}\to\text{epi}}$ and $\mathbf{z}_{\text{epi}\to\text{para}}$ through separate instances of the axial transformer. The resulting representations are concatenated and mean-pooled to obtain $\mathbf{z}_{\text{agg}}$.

###### E.3.4 Per-Residue Feature Integration and Transformer

After processing, pairwise features are broadcast back to residue positions and concatenated with single-residue embeddings:

$$\mathbf{h}'_i = [\mathbf{h}_i; \mathbf{z}_i] \in \mathbb{R}^{d_h + d_{z'}} \tag{8}$$

where $\mathbf{z}_i$ denotes the pairwise features associated with residue $i$.

The combined per-residue representations are then processed by a transformer encoder with 2 layers and 2 attention heads. This enables cross-residue information flow, allowing the model to capture cooperative effects between interface positions. The output is aggregated via masked mean pooling over non-padded positions:

$$\mathbf{h}_{\text{pooled}} = \frac{1}{L}\sum_{i=1}^{L} \mathbf{h}''_i \tag{9}$$

where $L = n_{\text{para}} + n_{\text{epi}}$.

###### E.3.5 Classification Head

Global confidence features $\tilde{\mathbf{c}}$ are concatenated to the pooled representation alongside the aggregated pairwise features. The final classifier consists of a two-layer MLP followed by a linear projection:

$$\hat{\mathbf{y}} = \mathbf{W}_{\text{out}} \cdot \text{Dropout}\left(\text{ReLU}\left(\mathbf{W}_{\text{clf}}\left[\text{pool}\left(\text{Transformer}\left([\mathbf{h}; \mathbf{z}_{\text{agg}}]\right)\right); \tilde{\mathbf{c}}\right] + \mathbf{b}_{\text{clf}}\right)\right) + \mathbf{b}_{\text{out}} \tag{10}$$

where the hidden dimension is 16 and output dimension is 2 (binding vs. non-binding classes).

#### F ML with SEPIA: Supporting Analysis

##### F.1 Cross-validation Preparation

We created 5-fold cross-validation splits of SEPIA data for *in silico* ML experiments, designed to prevent information leakage. We used a graph-based approach to identify closely related sequence clusters of VHHs, SEPs, and antigens, and accounted for pairings of VHH–Ag and VHH–SEP where the VHH is shared but the cognate differs.

First, we collected all unique on-target VHH–SEP pairs (binding and non-binding) from Round 1 and Round 2 as well as all unique on-target VHH–Ag pairs from SAbDab-nano. VHH sequences are clustered using `mmseqs2` [24] with 95% identity and 90% coverage; the same is done to cluster sequences of SEPs and of antigens from SAbDab-nano. These clusters each form a cluster node in our graph. Every unique sequence of VHH, SEP, and antigen is also a node, and they are all connected to their respective `mmseqs2` cluster nodes with an edge in the graph. Then, for all VHH–SEP

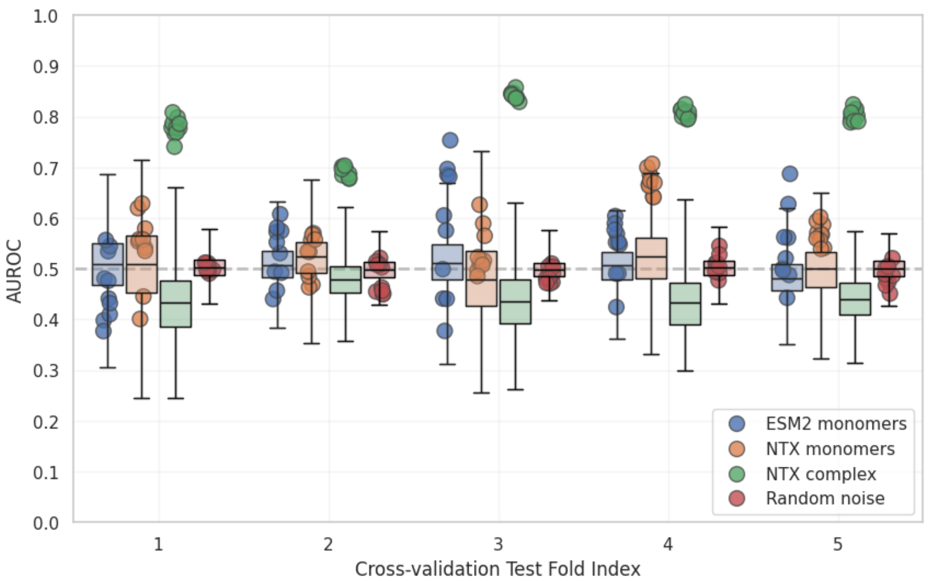

**Figure S20: Label ablation study**: AUROC test set performance from 5-fold cross-validation of GBM classifiers fit to real VHH–SEP pair hit/miss labels (circles) versus GBM classifiers fit to random permutations of those labels that form a null distribution (box plots). Four different representations of the VHH–SEP pairs are compared: ESM for a sequence-only representation (blue), structure-aware NTX embeddings of non-interacting VHH and SEP monomers (orange), structure-aware NTX embeddings of the interacting VHH–SEP complex (green), and random Gaussian noise features (red). Ten resamplings were run for each test set, each with 100 random permutations of the training set labels (before train/validation split), amounting to 1000 null distribution data points shown for each test set. For early stopping, each GBM classifier made an 80/20 train/validation split on its respective training data. Box plots of the permuted data show sampled null distributions with the whiskers representing minimum and maximum values. The random baseline of AUROC = 0.5 is shown as a dash line for reference.

or VHH–Ag pairs, we draw an edge between that pair's VHH node and SEP/Ag node. Finally, we search the global graph to find all disjoint subgraphs. Nodes in each disjoint subgraph are marked as belonging to an atomic "group" for cross-validation that cannot be subdivided across train/test splits. Each group represents VHH–SEP/VHH–Ag pairs that may leak information indirectly due to sequence similarity or via their cognate identity, which we intend to practically minimize.

To form 5-fold train/test splits, we apply `StratifiedGroupKFold` from `scikit-learn` [20] to approximately balance bind/no-bind labels as well as volumes of each test set. Off-target data points from the SAbDab-nano decoy data set (Section D.1) that do not share a VHH or antigen with any of the on-target data points are not assigned a test fold index, allowing them to appear as additional negative data in training sets but not affecting test set evaluation. These off-target data points are confirmed to also be less than 95% sequence similar to test sets. The resulting train/test splits ensure that data points in each training set are 1) less than 95% sequence similar to any in the test set and 2) have not appeared as a member of any VHH–SEP/VHH–Ag pairs in the test set.

#### F.2 Label Ablation Study

We perform a null hypothesis test via label ablation to verify that SEPIA pseudo-structures have a statistically significant correlation with AlphaSeq affinity. The goal is to compare fitting ML models to real AlphaSeq labels versus random noise, ideally observing that model performance on the real labels is much better as an indication of learning a real, informative signal present in the pseudo-structure data. Results of this experiment are shown in Figure S20.

To capture the relationship between pseudo-structure feature representation and affinity label, we consider 4 featurizations of SEPIA pseudo-structures: 1) ESM2 [25] for sequence-only, monomer, structure-unaware 2) NTX monomer embeddings for structure-aware but not complex-aware 3) NTX complex embeddings for structure- and complex-aware 4) Gaussian noise for a random baseline. ESM2 embeddings were computed by embedding each VHH and antigen monomer sequence separately, mean-pooling over length, and concatenating. Gaussian noise features were sampled from $\mathcal{N}(0, 1)$ (1,000 features per data point). NTX monomer embeddings, $Z_{\text{monomer}}$, are produced as described in

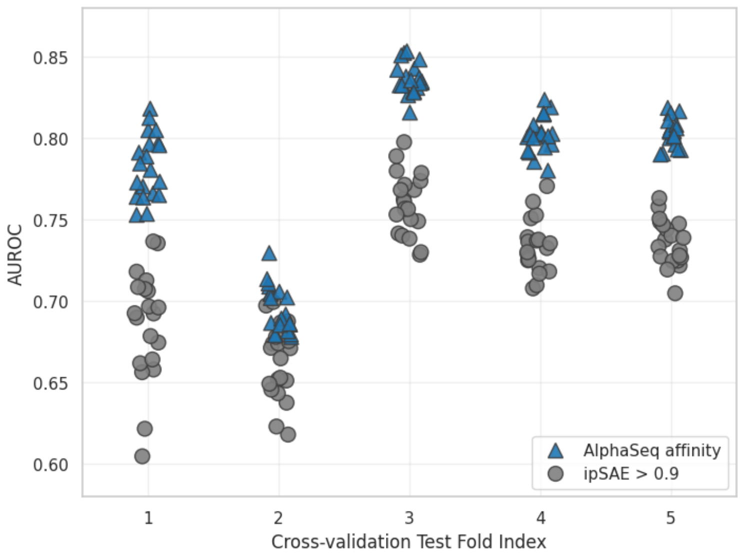

**Figure S21: Pseudo-labeling with high-confidence ipSAE**: AUROC test set performance on true AlphaSeq labels for GBM classifiers fit with true AlphaSeq labels (blue) versus fit with high-confidence ipSAE > 0.9 binarized pseudo-labels (gray); 5-fold cross-validation with 20× resamplings.

Section E.2.4, where NTX embeddings are computed for the isolated VHH and antigen in their holo conformation from the complex (i.e., same geometry but non-interacting) and concatenated similar to the ESM2 embeddings. The NTX complex embeddings, $Z$, also described in Section E.2.4, are the summation of confidence weighted differences of NTX embeddings, focusing on the interfacial residues.

Using the 5-fold cross-validation splits of SEPIA Round 1 data described in Section F.1, we generate a null label distribution for each training set by randomly permuting the true AlphaSeq bind/no-bind labels; test set true labels are left intact for evaluation. Within each train/test split and for each of the 4 featurizations, we fit a histogram gradient boosting machine (GBM) model on both the true AlphaSeq labels and on the randomly permuted labels. Each GBM model uses a stratified 80/20 train/validation split for early stopping. We perform 10× resamplings of the cross-validation to account for variance in the train/validation splits, as well as 100× label permutations per resampling, such that in total there are 1,000 GBM model permutation fits for the null distribution. The performance of the GBM models is evaluated as AUROC on the true test set labels. This allows for a calculation of $p$-value as the fraction of times where a GBM model fit to permuted labels has an AUROC greater than any GBM model fit to the true labels.

First, we note that all featurizations yield a null distribution encompassing AUROC=0.5, as expected, but with somewhat different amounts of variance and bias. Notably, the NTX complex null distribution appears biased below AUROC=0.5, which is apparently due an interplay between the class imbalance and latent space organization of real positive labels. When fit to the real labels, we see the GBMs fit to Gaussian noise all fall within the minimum/maximum bounds of the corresponding null distribution, implying there is not significant association between Gaussian noise and the AlphaSeq labels, as expected. GBMs fit using ESM2 features (monomer, sequence-only) are also observed to not have significant association to the AlphaSeq affinity labels, all but a few falling within the bounds of the null distribution, implying that VHH–SEP binding is not trivially encoded by the VHH or SEP amino acid sequence. GBMs fit to the NTX monomer embeddings display occasional performance beyond the null distribution for fold 4, but overall they are largely overlapping, which implies that VHH–SEP binding is also not trivially encoded in monomer structure alone. Finally, GBMs fit to the NTX complex embeddings are found to all perform well beyond the limits of the null distribution across all cross-validation folds, each amounting to a $p$-value < 0.001, given the 1,000 permutations.

We conclude that SEPIA pseudo-structures contain meaningful information about AlphaSeq binding affinity that ML models can learn, on the condition that the featurization of the pseudo-structures is aware of the full VHH–Ag complex structure and amino acid sequences, as in the NTX complex embeddings.

##### F.3 Pseudo-labeling Study

We investigated how well a GBM classifier performs when trained on high-confidence ipSAE pseudo-labels (binarized at ipSAE > 0.9) in comparison to being trained on the *in vitro* AlphaSeq affinity labels. We carried out a 5-fold

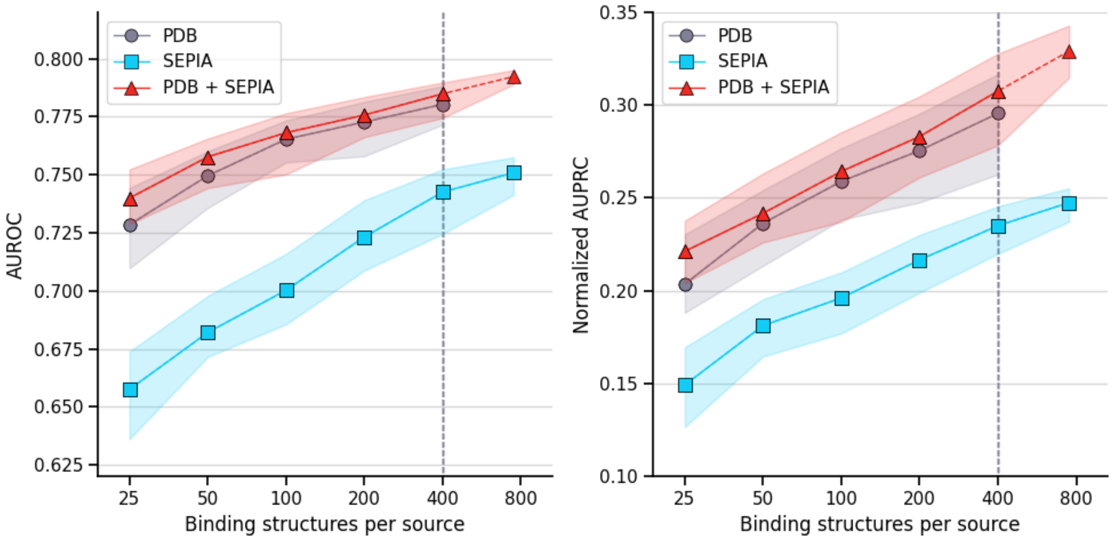

**Figure S22: Cross-validation learning curves**: Model performance at varying training data volumes at the task of classifying if a structure is a real or decoy VHH-antigen complex, evaluated on the decoy dataset. MLP classifiers are trained with the PDB-derived decoy data (gray), the SEPIA dataset (blue), and their union (red). The AUROC (left) and normalized AUPRC [26] (right) are computed globally over all test set data points (i.e., ranking predictions across all VHH-antigens pairs, not subset or averaged by VHH or antigen) by concatenating predictions on the 5 test sets from each single cross-validation pass. The dashed vertical line at x=400 represents the effective limit of PDB data in this cross-validation experiment.

cross-validation similar to what is described in Section F.2 using Round 1 SEPIA data and 20× resampling. NTX complex embeddings are used as feature inputs for GBM classifiers trained with early stopping on 80/20 stratified train/validation splits.

Results are shown in Figure S21. The ipSAE-derived pseudo-label models exhibit learning a signal, as is expected knowing that ipSAE can often correlate with binding. However, models trained on AlphaSeq labels visibly outperform models trained on the pseudo-labels.

##### F.4 Decoy Detection Learning Curves

We evaluated SEPIA data informational content and scaling behavior using the 5-fold cross-validation described in Section F.1, with the PDB-derived decoy dataset (Section D.1) as test sets. Within this cross-validation, training on decoys is effectively capped at 400 binding structures, having exhausted all available VHH–antigen structures. SEPIA data is analogously capped at 750 binding pseudo-structures. We compared three training conditions: (1) PDB-derived decoys only (PDB), (2) SEPIA Rounds 1 and 2 only (SEPIA), and (3) a 1:1 mixture (PDB + SEPIA), evaluating over sequential doublings of training set size with 100 random stratified resamplings at each volume and a fixed 1:10 binding/non-binding ratio.

For each resampling, a simple MLP classifier was fit using the same hyperparameters across all conditions: a 500-dimensional z-score normalized NTX embedding input (Section E.2.4), a single 10-dimensional hidden layer with sigmoid activation, binary cross-entropy loss, Adam optimizer (lr=$10^{-3}$, batch size 200, L2 weight decay 0.1, stratified mini-batch sampling), light Gaussian input noise of $\mathcal{N}(0, 0.1)$ during training, and early stopping with 20 epochs patience on a 20% validation split. Performance metrics were computed by concatenating test set predictions across all 5 folds for each of the 100 resamplings.

As shown in Figure S22, training on SEPIA alone yields continual improvement across all metrics (despite being ostensibly out-of-distribution for the decoy detection task), eventually reaching performance comparable to PDB-derived decoys at 50 binding data points and with a similar upward slope, suggestive of further gains possible with more data. The mixed PDB + SEPIA curve tracks the slope of the PDB-only curve but scales beyond its effective limit, achieving higher performance at the combined effective limit of 1,150 binding structures. These observed results depend on model choices and the bias-variance tradeoff; we have aimed to balance model complexity and regularization to observe the effects of data scaling. More exhaustive hyperparameter sweeps could likely yield higher absolute performance.

**Table S13:** Classification performance metrics on SEPIA data (mean ± std).

| Evaluation | Metric | ABACUS | ipSAE |
|---|---|---|---|
| Global | AUROC | $\mathbf{0.89 \pm 0.02}$ | $0.86 \pm 0.02$ |
| | AUPRC | $\mathbf{0.27 \pm 0.04}$ | $0.19 \pm 0.04$ |
| Per-VHH Aggregated | AUROC | $\mathbf{0.80 \pm 0.21}$ | $0.78 \pm 0.22$ |
| | AUPRC | $\mathbf{0.27 \pm 0.24}$ | $0.22 \pm 0.22$ |

#### F.5 SEPIA In-distribution Performance Evaluation

In this section we investigate the ABACUS's performance on SEPIA itself. We evaluated model performance using classification metrics (AUROC, AUPRC), enrichment metrics at various selection thresholds, and per-VHH aggregated statistics across 5-fold cross-val idation.

Table S13 reports global and per-VHH aggregated classification metrics (mean ± std across 5-fold cross-validation) and Figure S23 plots precision, enrichment, and recall across different quantiles.

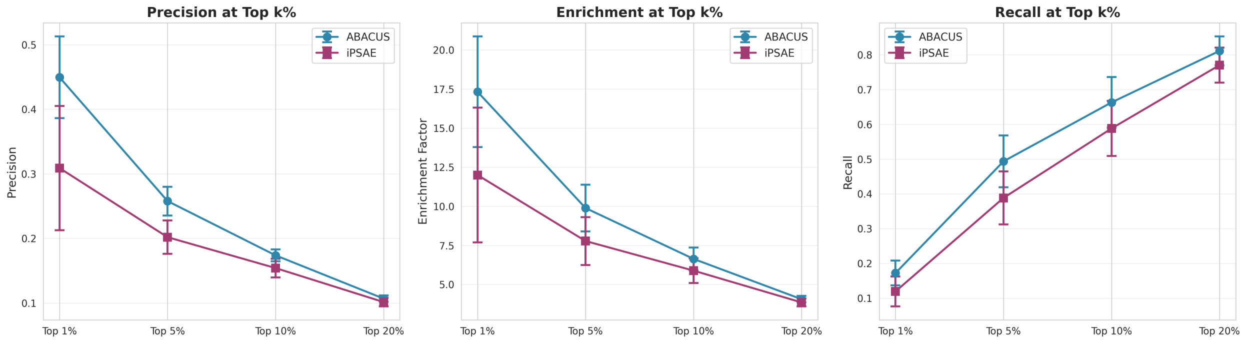

**Figure S23: Enrichment metrics on SEPIA data at different selection thresholds.** Line plots show mean performance across cross-validation folds with error bars representing standard deviation. (Left) Precision@k, (Middle) Enrichment@k, (Right) Recall@k.

#### F.6 Combining SEPIA and Decoys in ABACUS Training Data

We investigated the effect of combining the PDB-derived decoy data with SEPIA data for ABACUS training, as well as training on decoy data alone. Including decoys to our SEPIA data generally improved the model, however global AUROC and the Enrichment@k AUC was observed to be slightly less than training on SEPIA data alone. (Figure S24, Figure S25). With the exception of antigen-averaged AUPRC, training on decoys alone performs worse than training on SEPIA alone. We zeroed out the confidence score features when training on decoy dataset alone, as this improved performance, likely because decoy structures overlap with Boltz-2's training data, making the confidence–binding relationship in this dataset misleading for novel targets. However, when combined with SEPIA, we did not see a benefit to zeroing out the confidence scores for the decoy dataset.

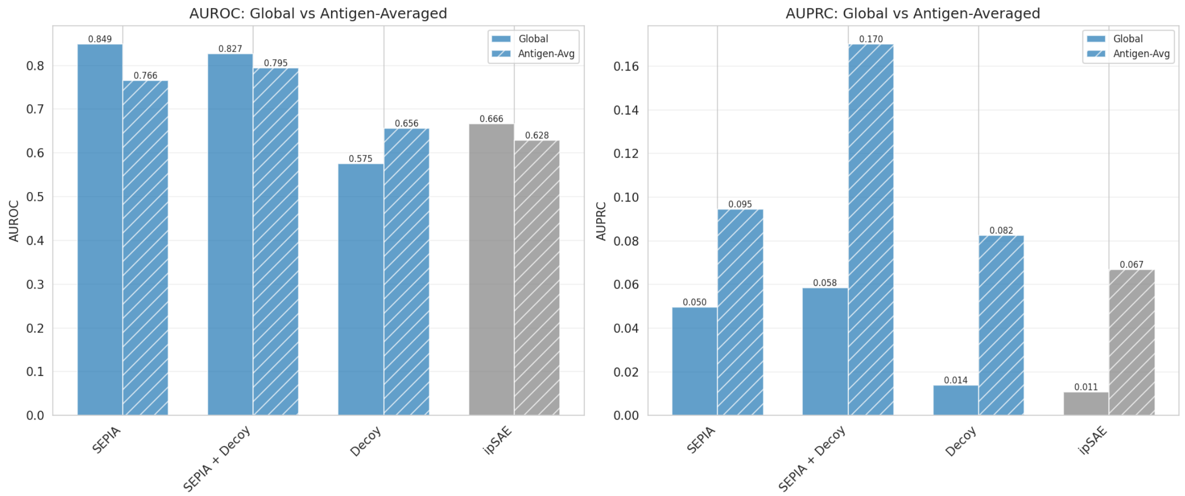

**Figure S24:** ***De novo*** **binding classification performance across training data sources.** Comparison of ipSAE with models trained on SEPIA data alone, decoy data alone, or both combined.

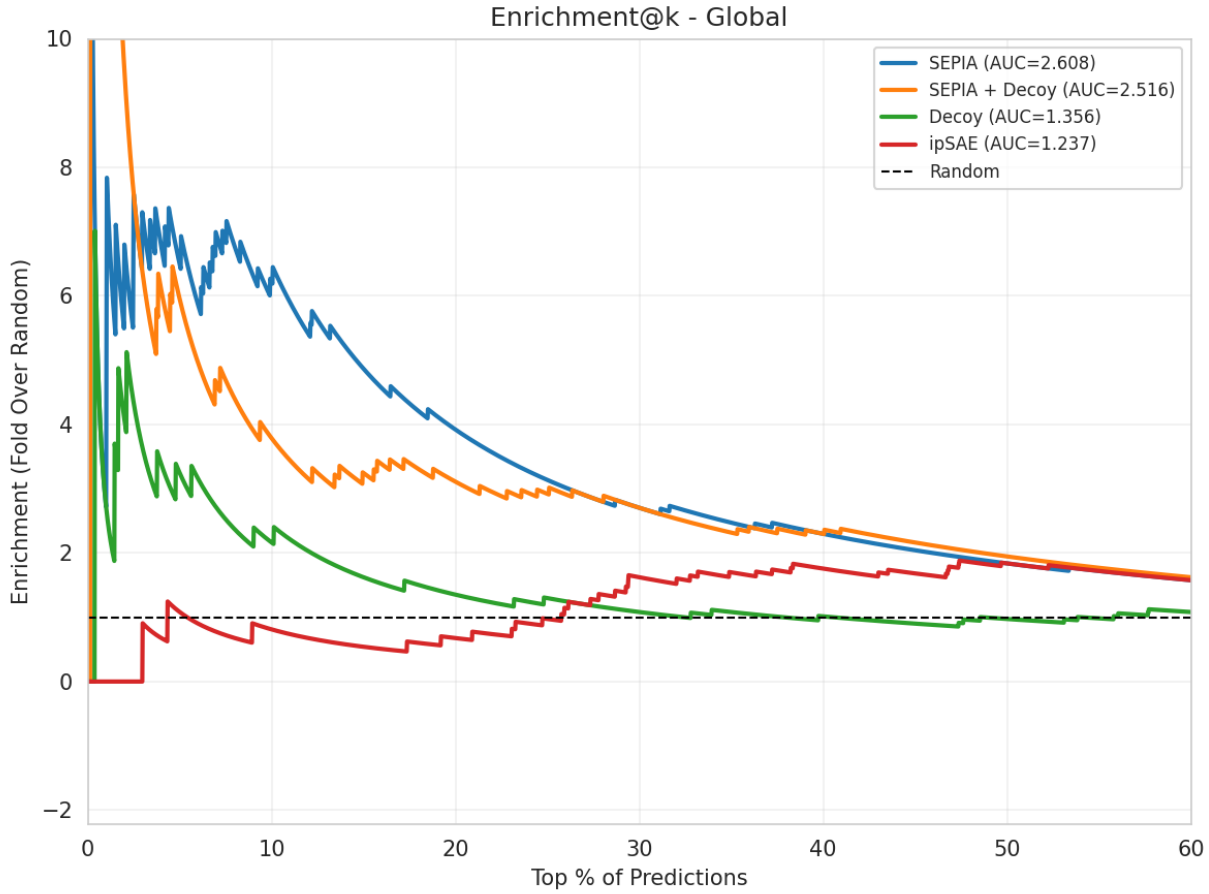

**Figure S25: Enrichment performance comparison across training data sources.** Enrichment@$k$ curves for models trained on SEPIA data, PDB-derived decoy data, or both. By enrichment AUC, training on SEPIA alone performs best, followed closely by SEPIA combined with decoys, even though the latter exhibits higher enrichment at the very top percentiles. Training only on decoys outperforms ipSAE yet performs considerably worse than the models trained on SEPIA with/without decoy data.

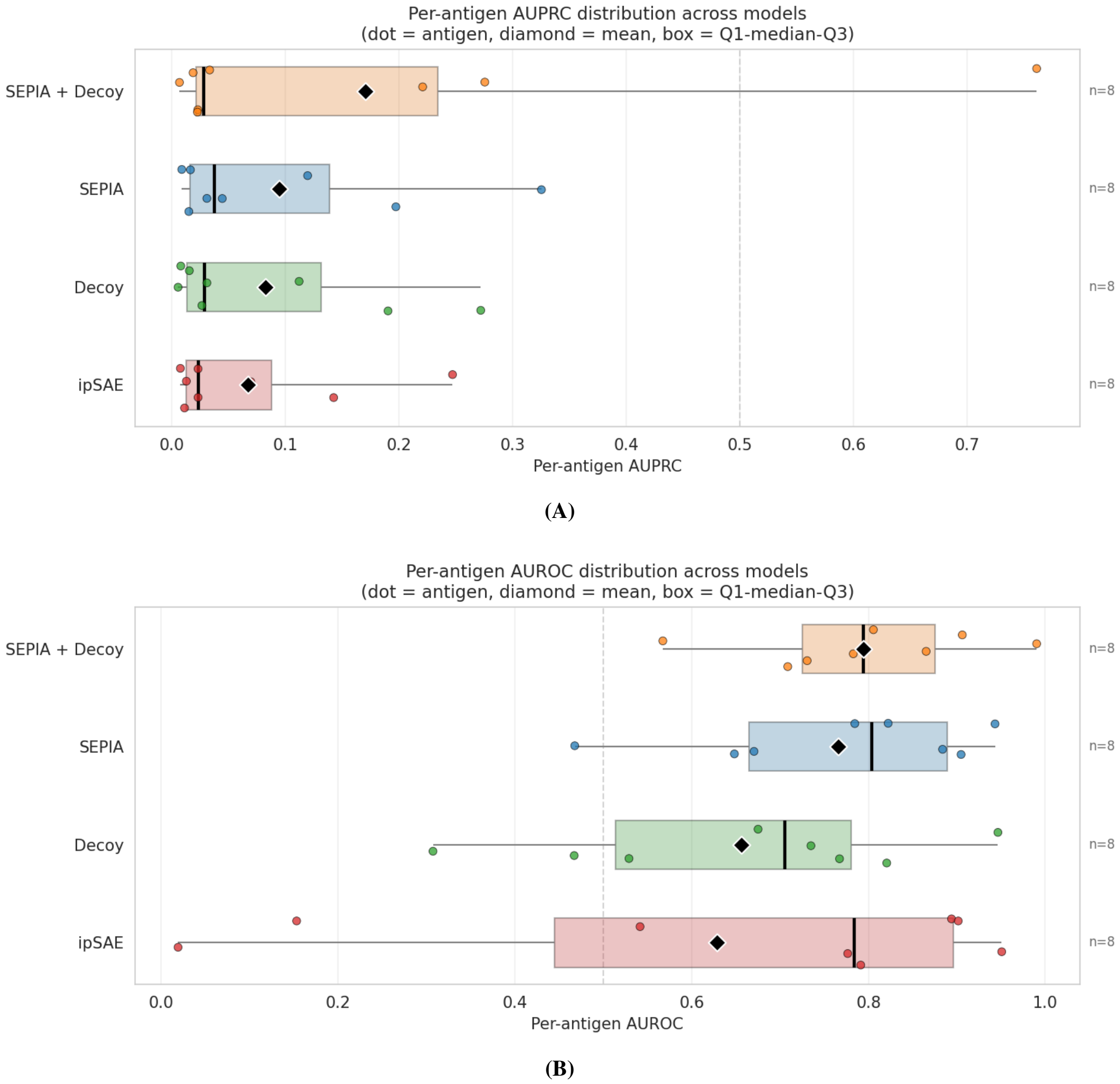

**Figure S26: Per-antigen classification performance across training data sources.** (A) AUPRC and (B) AUROC broken down by antigen for models trained on SEPIA data, PDB-derived decoy data, or both.
